# Nanoscale Friction of Biomimetic Hair Surfaces

**DOI:** 10.1101/2022.09.29.510078

**Authors:** Erik Weiand, James P. Ewen, Yuri Roiter, Peter H. Koenig, Steven H. Page, Francisco Rodriguez-Ropero, Stefano Angioletti-Uberti, Daniele Dini

**Affiliations:** Department of Mechanical Engineering, Imperial College London, South Kensington Campus, SW7 2AZ London, U.K.; Institute of Molecular Science and Engineering, Imperial College London, South Kensington Campus, SW7 2AZ London, U.K.; Thomas Young Centre for the Theory and Simulation of Materials, Imperial College London, South Kensington Campus, SW7 2AZ London, U.K.; Corporate Functions Analytical and Data & Modeling Sciences, Mason Business Center, The Procter and Gamble Company, Mason, 45040 Ohio, U.S.A.; Department of Materials, Imperial College London, South Kensington Campus, SW7 2AZ London, U.K.

## Abstract

We investigate the nanoscale friction between biomimetic hair surfaces using chemical colloidal probe atomic force microscopy experiments and nonequilibrium molecular dynamics simulations. In the experiments, friction is measured between water-lubricated silica surfaces functionalised with monolayers of either octadecyl or sulfonate groups, which are representative of the surfaces of virgin and ultimately bleached hair, respectively. In the simulations, friction is monitored between coarse-grained model hair surfaces with different levels of chemical damage, where different fractions of grafted lipid molecules are randomly replaced with sulfonate groups. The sliding velocity dependence of friction can be described using an extended stress-augmented thermally activation model. As the damage level increases, the friction generally increases, but its sliding velocity-dependence decreases. At low sliding speeds, which are closer to those encountered physiologically and experimentally, we observe a monotonic increase of friction with the damage ratio, which is consistent with our new experiments using biomimetic surfaces and previous ones using real hair. This observation demonstrates that modified surface chemistry, rather than roughness changes or subsurface damage, control the increase in nanoscale friction of damaged hair. We expect the experimental and computational model surfaces proposed here to be useful to screen the tribological performance of hair care formulations.

## Introduction

A detailed understanding of the tribology of human hair is essential to develop new shampoos and conditioners that leave hair untangled and feeling smooth.^1^ In particular, maintaining low friction between hairs is important for satisfactory sensory perception during touching, brushing and combing.^2–7^ Consequently, the kinetic friction of hair has been studied using a wide range of experimental techniques from the macroscale to the nanoscale.^8–30^ The friction between nanoscale tips and single hairs, ^10–14,18,22,26^ as well as crossed hair-hair contacts,^23–25^ have been investigated using atomic force microscopy (AFM) and high-load nanotribometers. ^27^ Understanding the friction between hairs is crucial since this dominates the overall resistance felt during combing and brushing. ^25^

Friction forces on hairs are anisotropic due to the overlapping nature of the outer cuticle cells.^9^ Friction is much higher when hairs are rubbed in the tip-to-root direction, where the cuticles lock together, than in the root-to-tip direction, where they are able to slide over one another more easily.^9^ A monolayer of 18-methyleicosanoic acid (18-MEA) is covalently bonded to the protein layer below, mostly through thioester bonds with cysteine residues in the cuticle of the hair. This protective 18-MEA monolayer is commonly known as the fatty acid layer (F-layer). The F-layer makes hair surface hydrophobic and also provides low friction.^12,29^ Previous AFM experiments have consistently found an increase of the coefficient of friction (CoF) for bleached hair compared to virgin hair.^10,12–14,22,24^ During bleaching, the 18-MEA layer is partially removed by oxidation of the cysteine, which leads to the formation of negatively charged cysteic acid groups, making the hair surface more hydrophilic.^19,31–33^ From experiments in humid air environments (relative humidity, RH ≈50%), capillary condensation has been proposed as a possible mechanism for the increase in adhesion and friction from bleaching hair. ^12^ Indeed, capillary condensation is known to increase friction on hydrophilic surfaces more than hydrophobic ones. ^34^ However, more recent investigations showed that even in dry environments (RH ≈4%) such increases in the CoF are observed for bleached and chemically damaged hair. ^22^ The relative increase in CoFs on chemically damaged hair was found to be largest for measurements with a nanoscale AFM tip, as compared to microscopic and macroscopic methods, where cuticle edge effects and probe size can perturb the friction signals. ^15^ This motivates an investigation of hair friction phenomena at the smallest scales, where effects due to changes in surface chemistry can be isolated from topographical effects.

Environmental factors also affect hair friction. For example, hairs undergo swelling when soaked in water, ^16^ which has been found to lead to increased friction forces. It has been suggested that the hair surface softens in water, leading to larger contact areas and thus friction. ^22,24^ The application of cationic surfactant-based hair conditioners on damaged hair can partially recover the hydrophobic character of virgin hair. ^29^ Conditioners also lower the CoF compared to chemically damaged hair^14^ by restoring the boundary layer that was partially lost when 18-MEA molecules were removed.^24^

As with most natural materials, the structure and friction of hair shows considerable variability between individuals and populations. ^13^ The development of synthetic biomimetic surfaces with repeatable properties would therefore be useful for the screening of the tribological performance of different hair care formulations.^1^ Artificial hair mimics have been constructed to reproduce the microscale roughness features of overlapping cuticles^35^ and the F-layer has been represented by a gold AFM tip coated with a thiol monolayer. ^18^ However, the friction of biomimetic surfaces for chemically damaged hair has not yet been investigated. The use of atomically-smooth, biomimetic surfaces in nanotribology tests allows for reduced variability compared to real hair and for the detailed investigation of the effects of surface chemistry and treatments, while eliminating the influence of microscale roughness, which otherwise leads to friction anisotropy. ^9^

Confined nonequilibrium molecular dynamics (NEMD) simulations have been used to study the friction behaviour of a wide range of systems. ^36^ Surfactant monolayers adsorbed on solid surfaces are one of the of the most widely studied systems with NEMD simulations due to their importance in engineering and biological systems. These studies have investigated a wide range of applications from organic friction modifier (OFM) lubricant additives^37,38^ to synovial joints.^39,40^ While most of these studies have employed all-atom force fields,^36^ the large, heterogenous surfaces and macromolecules commonly encountered in biological systems are more suited to the use of coarse-grained representations.^41^ Coarse-grained force fields have already been used to study, for example, the intermonolayer friction of lipid bilayer membranes^42,43^ the friction of protein translocation through nanopores,^44^ and the adsorption and desorption of polymers on heterogeneous substrates that are representative of the surface of the hair under shear flow.^45^

In this study, we use NEMD simulations and chemical colloidal probe (CCP) AFM to study the kinetic friction between model hair surfaces. In the AFM experiments, we study the friction of water-lubricated biomimetic surfaces that are representative of virgin and ultimately bleached hair. In the NEMD simulations, we employ our recently developed and validated coarse-grained (CG) model of the surfaces of virgin and bleached hair^33^ within the *MARTINI* framework.^46^ Both dry and wet contacts are considered under a wide range of sliding velocities. Both the experimental and simulation frameworks will serve as a useful benchmark for the high-throughput screening of the tribological performance of potential hair care formulations. ^1^

## Methodology

### MD System Setup

Coarse-grained molecular models of hair surfaces at different degrees of damage, as introduced in our previous study,^33^ are used to describe the contact interface. Here, the *MARTINI 2.0* force field^46,48^ is used in conjunction with a polarizable water model.^49^ The polarizable water model was selected because electrostatic screening is believed to play an important role for interactions between the damaged (charged) model hair surfaces. This model employs a 4:1 mapping for non-hydrogen atoms to coarse-grained beads. Full details of the parameters chosen can be found in our previous study. ^33^ The systems were constructed using the Moltemplate software.^50^

In the *MARTINI* model, a shifted Lennard-Jones (LJ) potential^51^ is used to describe the non-bonded interactions. Switching to zero is performed between the cut-off radius *r*_LJ,cut_ = 0.9 and *r*_LJ,shift_ = 1.2 nm.^46^ Coulomb potentials are added for interactions between charged beads. Long-range electrostatic interactions are considered using the Yeh-Berkowitz slab implementation^52^ of the particle-particle particle-mesh (PPPM) method.^53^ A Coulombic switching radius of *r*_C,cut_ = 1.2 nm and a relative energy tolerance of 10^−5^ are applied. Bonds and angles are treated using weak harmonic potentials as in the original *MARTINI* framework. ^46^ For wet systems, bonds between polarizable water beads of a single water unit are constrained using the SHAKE^54^ algorithm.

Fig. 1 shows a representative example of a dry and wet system (fully-functionalised) during compression and sliding. A graphene sheet is used as an impenetrable substrate for grafting the lipid monolayer at a separation distance *d*_graft_ = 0.65 nm. This separation distance reproduces the experimentally-measured thickness of the F-layer. ^33,55^ Harmonic bonds are added between the coarse-grained carbon chain beads (C_1_).^46^ Deformation of the graphene beads in the *z* (surface-normal) direction is possible, but they are constrained in the *x* and *y* directions. As in our previous study,^33^ the lipid grafting positions are independent from the graphene beads at a nominal grafting distance of *d*_graft_ = 0.65 nm in a hexagonal lattice (surface coverage = 2.7 molecules nm^−2^), which is achieved using ghost beads in the same plane as the hexagonal sheet. The ghost beads are coupled to the motion of the graphene sheet during compression and shearing. The hexagonal arrangement of the molecules is motivated by experimental observations. ^56^

**Figure 1:**
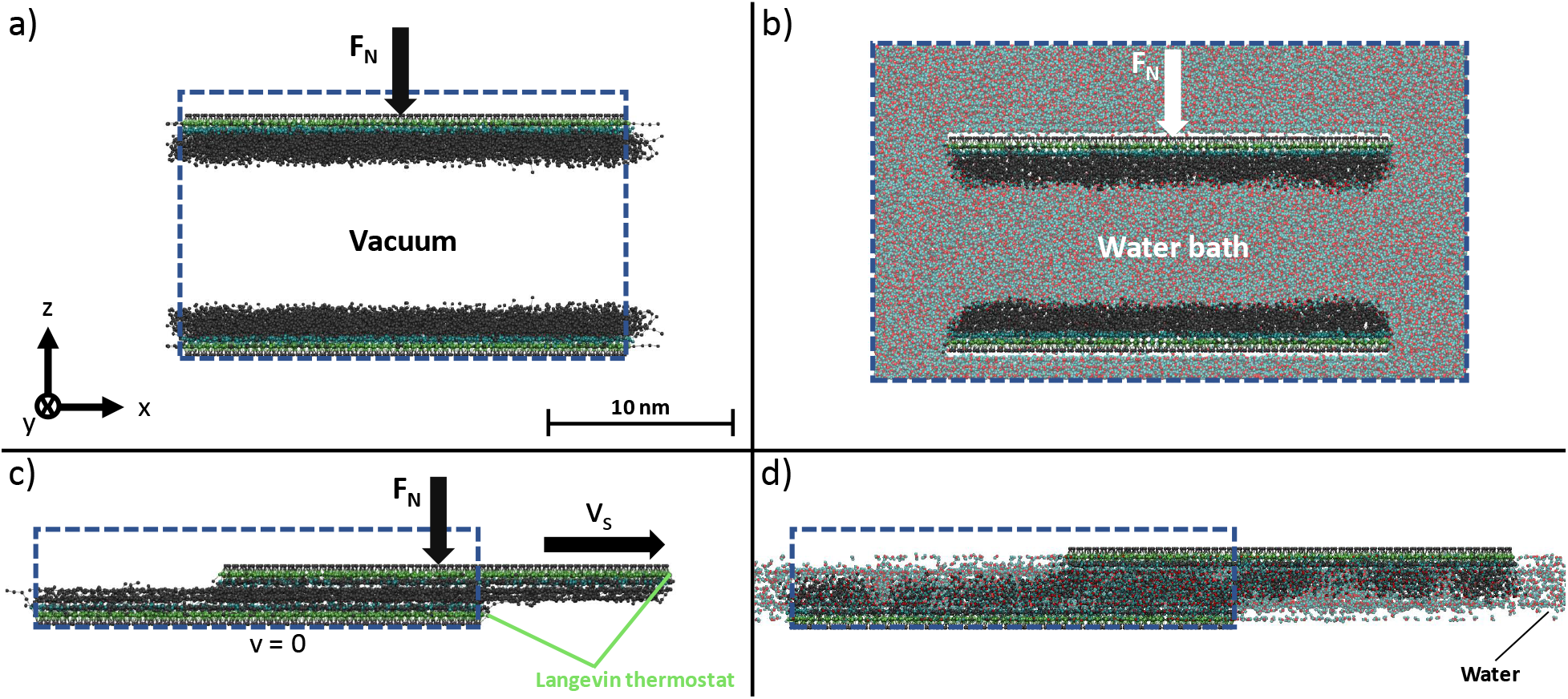
CG-MD configurations of squeeze-out systems in a) dry, b) wet contacts and sliding contacts for c) dry and d) wet contacts. Sliding contacts are shown in the unwrapped state as opposed to the periodic, wrapped bead coordinates to indicate the relative motion. Beads are coloured by type: C_1_ (gray), C_5_ (cyan), P_5_ (lime) and water (central red, satellite cyan opaque). Rendered with VMD.^47^

The surface dimensions are 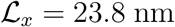 and 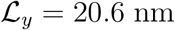. Given that the thickness of a human hair is typically 50-100 μm and each cuticle cell is approximately 5-10 μm long,^19^ this represents a small patch of a single cuticle. We tested the effect of surface size on the convergence of thermodynamic and friction properties in preliminary simulations. No statistically significant differences were found between the final surface size and simulations with an increase in edge length of 50% in *x* and *y*. We suggest that smaller surfaces should not be used for this purpose to ensure resolving characteristic damage patterns, particularly at low degrees of hair damage, where only a few damage islands exist on the surface.

We consider an idealized contact between the cuticle F-layers of two hair fibers. The contact is treated as atomically flat, which is an adequate simplification at the length scales investigated here (curvature = 0.02° at 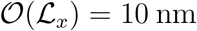 for *d*_hair_ = 50 μm). The lipids are representative of 18-MEA chains covalently bonded to the underlying protein layers via thioester bonds to cysteine groups. In the *MARTINI* framework,^46,48^ this consists of a surface-grafted P_5_ bead bonded to a C_5_ bead (representative of the thioester and amine/carboxyl groups in cysteine) bonded to five C_1_ beads (representative of the eicosane chain). ^57^ The oxidised cysteic acid groups formed by damage from chemical treatments or bleaching are represented by sulfonate groups that are covalently bound to the underlying protein. ^32^ The sulfonate groups are represented by Q_a_ beads (replacing the neutral C_5_ bead) with a charge of −1.^58^

The surface damage is quantified by the number damage ratio χ_N_, i.e., the ratio of the number of eicosane chains randomly replaced by sulfonate groups to the number of eicosane chains in a pristine monolayer. ^33^ We consider such fully-functionalised monolayers (χ_N_ = 0), as well as those we showed previously to be representative of the surface of virgin hair (χ_N_ = 0.25) and medium bleached hair (χ_N_ = 0.85),^33^ as well as ultimately bleached (χ_N_ = 1.0) and other intermediate degrees of damage. The degree of damage was calibrated to the different hair types through comparison to experimental contact angle data using water and *n*-hexadecane.^33^ For the damaged surfaces, the required number of sodium counter ions (Na^+^) are added to the systems to maintain charge neutrality. The Na^+^ cations and hydration shells (three water molecules) are represented by the Q_d_ beads with a charge of +1.^48^ The contacts studied in the NEMD simulations consist of two surfaces with an equal degree of damage. Excluding the screening effect of the counterions, the damaged surfaces carry a surface charge density reaching from *ρ_q_* = −11.0 *μ*C cm^−2^ to *ρ_q_* = −37.2 μC cm^−2^ for virgin and medium bleached hair, respectively. These values are somewhat lower than those from previous experimental measurements of the surface charge of virgin (−1.5 *μ*C cm^−2^) and bleached (−10.0 *μ*C cm^−2^) hair.^59^

### MD Simulation Details

Classical MD simulations are performed using the LAMMPS software. ^60^ The velocity-Verlet^61^ integration scheme is used with a timestep of 5 fs. First, the systems are energy minimized using the conjugate gradient algorithm, before being equilibrated at *T* = 298 K with a global Nosé-Hoover thermostat.^62,63^ During this phase, the position of the graphene sheets is fixed in the *z*-direction. Next, a constant normal force (*F_N_* = 4.9 nN) equivalent to a pressure of 10 MPa is added to all of the beads in the top graphene sheet (A = 490 nm^2^), while the beads in the bottom graphene sheet remain fixed. The chosen pressure is based on an estimate of the Hertz pressure in previous AFM experiments of friction in hair-hair contacts.^24,25^ Here, the applied load, *F_N_* ≈ 0.1 mN, and the Young’ modulus of virgin and chemically damaged hair were taken as *E* = 0.9 GPa or *E* = 0.5 GPa, which is representative of virgin and chemically damaged soaked hair, respectively.^64^ Further details are shown in the Supplementary Material (Fig. S1).

In the dry systems, the force is maintained until the average pressure reaches the target value. The dry systems are periodic in the *x* and *y* directions and finite in the z direction. In the wet systems, the normal force is maintained until the number of water molecules remaining in the contact and corresponding contact thickness reach a steady state value, to simulate the squeeze-out process at the target pressure. ^38^ For these squeeze-out simulations, a fully periodic system is used, as shown in Fig. 1 b). Water beads are initially distributed randomly in the contact using the Moltemplate software^50^ with additional reservoir space added around the contact in the *x* direction. No gap is added between the surfaces and the periodic boundary in the *y* direction. The number of water units in the system is fixed throughout the squeeze-out simulations. The initial surface-normal separation distance between the two graphene layers of the surfaces is set at *d* = 11.5 nm. Before the compression phase, the system is equilibrated at 300 K and atmospheric pressure (1 atm) is followed using a Parrinello-Rahman barostat acting in the *z*-direction.^65^ Once the system volume reached a steady state value, the dimensions were fixed again for the compression phase. For the sliding phase of the wet contacts, new systems are constructing containing the steady state number of water beads from the squeeze-out simulations and the additional space in the *x*-direction is removed. The simulation boxes for the dry and wet NEMD simulations are periodic in the *x* and *y* directions and finite in the *z* direction. These systems are energy minimised, equilibrated at 300 K, and then compressed to 10 MPa prior to the sliding phase. The chosen pressure for the MD simulations is consistent with that calculated using Hertz theory for two crossed hair fibres using measured values of the Young’s Modulus of virgin and chemically damaged hair^64^ and the load range used in previous AFM experiments of hair-hair friction (Fig. S1).^24,25^ The chosen pressure is also representative of the CCP AFM experiments using the biomimetic surfaces (Fig. S2).

For the sliding phase, at sustained normal force, the upper graphene surface is moved in the *x*-direction at a constant sliding velocity, *v_s_*, while to bottom slab remains stationary. Sliding velocities between 10^−2^ and 10^0^ m s^−1^ are considered, which are several orders of magnitude higher than typical velocities in AFM experiments using hair that typically operate in the 10^−6^ to 10^−3^ m s^−1^ range.^16,18,24^ The *v_s_* values considered in the NEMD simulations are, in fact, closer to realistic scenarios of hair manipulations such as combing^66,67^ or rubbing with a finger.^68^ The requirement to sample a sufficient fraction of each surface in contact at reasonable computation times imposes a lower limit on the sliding velocity.

During the sliding phase, thermostatting is applied only to the P_5_ beads of the hair surfaces in the direction lateral (*y*) to compression and sliding to act as a heat sink for thermal dissipation from frictional heating close to the sliding graphene sheets. This is a more physically realistic thermostatting strategy compared to thermostatting the entire system, which prevents thermal gradients from developing. ^69^ For this purpose, a Langevin thermostat^70^ is applied to the P_5_ beads at *T* = 298 K and a time relaxation constant *τ* = 0.1 ps. Production sliding simulations are run for at least 200 ns. At the lowest sliding speeds considered here (*v_s_* = 0.01 m s^−1^), simulations are run up to 500 ns to sample at least 5 nm of surface displacement.

#### Water Model Viscosity

Since the viscosity of fluids is important to their tribological behaviour of soft contacts, ^71^ we validated the pressure-viscosity response of the polarizable *MARTINI* water model against experimental data. Viscosity calculations with the *MARTINI* force field have been performed in few instances,^42,43,72–74^ but the viscosity of the polarizable *MARTINI* water models have not been previously reported. Here, we employ the Green-Kubo method^75,76^ to determine the dynamic viscosity *η* of polarizable *MARTINI* bulk water^49^ at ambient conditions (*T* = 298 K, *p* =1 atm) in the NVT ensemble after equilibration using a Nosé-Hoover thermostat.^62,63^ A value of *η* = 54.8 ±1.5 mPas is found, which underestimates the experimental value^77^ by ~ 39 %. The magnitude of this deviation is comparable to that of other atomistic water models, ^78^ such as SPC/E^79^ or TIP5P.^80^ The size of the simulation cell did not significantly affect the viscosity for the three tested box lengths 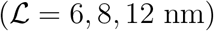 shown in the Supplementary Material (Fig. S3a). Furthermore, we evaluated the dynamic viscosity at system pressures of p = 0.1, 100, 200, 300 MPa using a Nosé-Hoover thermostat and barostat.^62,63^ The corresponding viscosity-pressure plots are shown in the Supplementary Material (Fig. S3b). Unlike for experimental water, the viscosity of the polarizable *MARTINI* model strongly increases with pressure. At high pressure, the model overestimates viscosity compared to experimental values. This is believed to result from the well-known attribute of *MARTINI* water to be prone to strong reordering, ^48^ thus transitioning to a solid-like behaviour at significantly lower bulk pressures than real water. We also observe this behaviour at elevated temperatures (*T* = 323 K), although shifted towards slightly higher bulk pressures. Thus, we conclude that the range of system pressures should be limited to *p* < 100 MPa to remain in a realistic viscosity regime. Such pressures are generally sufficient to match those observed experimentally inside the soft contacts found in biological systems. ^81^

### Experimental Details

We performed CCP AFM^82^ experiments in a liquid cell to study the friction of atomically-smooth hair mimic surfaces in a water-lubricated environment. Biomimetic hair surfaces were produced by silanizing silicon wafers (0.2 nm root-mean-square roughness) and 5 μm silica colloidal AFM probes with either octadecyl (C_18_) or sulfonate (SO_3_^−^) groups. The composition and properties of the monolayers were analysed using a spectroscopic ellipsometry, time-of-flight secondary ion mass spectrometry and contact angle measurements. Virgin hair was represented by a pristine C_18_-terminated self-assembled monolayer (SAM), while complete damage was represented by a SO_3_^−^—terminated SAM. The use of these biomimetic surfaces eliminates any damage-induced change in microscale surface topography^83^ or porosity^84^ from the friction measurements. These surfaces were similar to those proposed recently in our CG-MD wetting study on hair. ^33^

The CCP AFM friction experiments were performed using a Bruker (Billerica, Massachusetts) MultiMode 8 scanning probe microscope (SPM) and custom-ordered 5 μm colloidal silica NovaScan (Boone, Iowa) AFM probes attached to nominally 0.35 N m^−1^ Si_3_N_4_ cantilevers. Tipless reference cantilevers were used to calibrate the normal spring constant and microscopic geometrical calculations were used to calculate torsional/lateral spring constants. ^85^ CCP AFM friction studies were performed in deionised water medium. The typical scan length during measurements was 30-100 nm at 1 Hz frequency and normal loads 100-3000 nN. Separate CCP AFM measurements were made to quantify the repulsion in the SO_3_^−^-SO_3_^−^ systems (−20 nN) and about adhesion in the C_18_-C_18_ systems (+450 nN). This is consistent with previous AFM measurements that have shown reduced hair-hair adhesion following chemical damage^13^ or bleaching. ^24^

Using the Johnson-Kendall-Roberts (JKR)^86^ contact theory modified by Reedy^87^ for thin coatings with experimentally measured adhesion and elastic properties of the silica (substrate)^88^ and surfactant monolayers (coating),^89^ we estimate that the contact pressure in these experiments is 5-79 MPa. Further details on this calculation and additional estimation methods are given in the Supplementary Material (Fig. S2). Although the maximum estimated pressure is somewhat higher than in our NEMD simulations, we do not expect this to substantially affect our friction results. It is worth noting though that for previous AFM experiments using hair-hair contacts, the load had no effect on the results over a range of 10 to 1000 nN.^23^ High-load nanotribometer experiments of hair-hair contacts showed that the CoF decreased from 10 to 100 mN of normal load, at which point wear began to occur.^27^ Thus, we consider our chosen pressure in the NEMD simulations (10 MPa) to be representative of the current and previous experiments of nanoscale hair friction where no substantial wear occurs. We performed a subset of squeeze-out and NEMD simulations at higher pressures (20-50 MPa) to enable more direct comparison with the CCP AFM experiments.

## Results and Discussion

### NEMD Simulations of Dry Contacts

To help understand the molecular-scale origins of the friction behaviour, we first study the structure of the hair surfaces at different degrees of damage. Fig. 2 shows the surface-normal (*z*) mass density and through-film velocity profiles (*v_x_*) for dry contacts from the NEMD simulations. At low damage levels, the thickness of the F-layer on each surface is approximately 2 nm, which is consistent with previous experimental measurements. ^55^ At all damage levels, there is clear layering, which is strongest close to the sliding surfaces, but extends into the centre of the contact (*z* = 0 nm). The mass density profiles are insensitive to the sliding velocity, which is consistent with previous atomistic NEMD simulations of stearic acid monolayers on iron oxide surfaces. ^37^ The monolayers on the opposing surfaces are interdigitated with each other, as shown by the overlap in mass density profiles. There is a general increase in the amount of interdigitation with increasing damage ratio. The outer layers of the lipids move at the same velocity as the graphene surfaces to which they are grafted and the velocity changes linearly between the two extremes in the interdigitated region. This observation is also consistent with previous atomistic NEMD simulations of surfactant monolayers adsorbed on iron oxide surfaces. ^38^ There do not appear to be any clear velocity slip planes between the layers shown in the mass density profiles, which would appear as steep/discontinuous changes in the velocity profiles.

**Figure 2:**
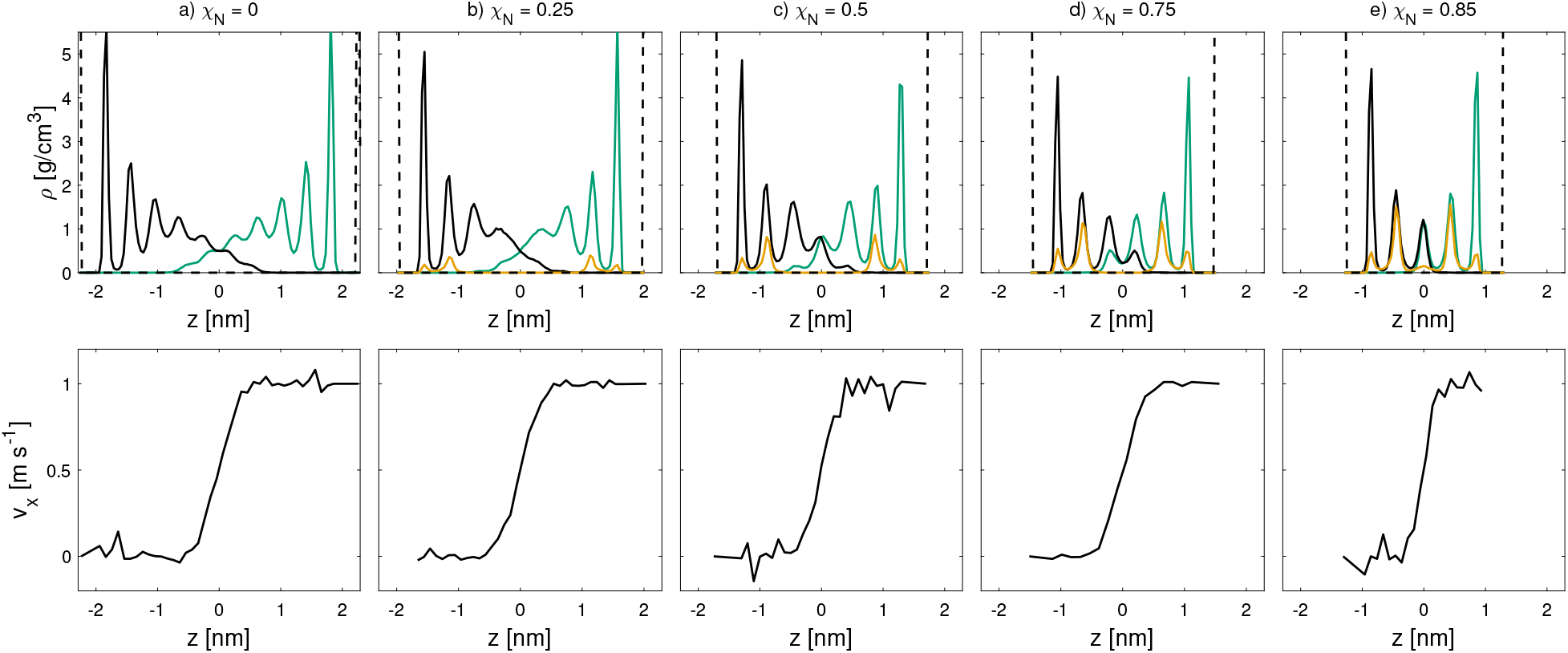
Dry contact monolayer mass density (top) and velocity (bottom) profiles at *v_s_* = 1 m s^−1^ and *σ* = 10 MPa for different damage ratios, χ_N_. The damage ratio increases from left to right. The densities of the lower surface (solid black), upper surface (solid green), sodium counterions (orange) and graphene sheets (black dashed) are shown.

The CoF (*μ*) is obtained using Amontons’ friction law by dividing the lateral force (*F_L_*) by the normal force (*F_N_*) acting on the outer layer of beads.^90^ Previous nanoscale AFM hair friction experiments have shown a linear relationship between *F_L_* and *F_N_*, with a negligible intercept. ^13,24^ The mean CoF between dry surfaces from the NEMD simulations is shown in Fig. 3 as a function of sliding velocity for the various levels of damage considered. At the lowest sliding velocity (*v_s_* = 0.01 m s^−1^) considered, the CoF increases monotonically with χ_N_, which is consistent with previous experimental measurements of virgin, chemically damaged and bleached hair. ^10,13,22^ At low sliding velocity, the NEMD simulations show a difference in CoF of almost two orders of magnitude between the virgin (χ_N_ = 0.25) and medium bleached hair (χ_N_ = 0.85). Experiments in a dry nitrogen atmosphere (RH ≈) showed that compared to virgin hair, the CoF was approximately three times higher for chemically damaged (KOH-treated) hair and twice as high for bleached hair.^22^ There are several differences between these previous experiments and the current simulations that may explain the larger difference between virgin and damaged hair in the simulations compared to the experiments. Firstly, the hair samples in the AFM experiments would have still contained some residual water molecules, despite the low relative humidity. Secondly, the experiments used silicon nitride tip-hair contacts, rather than hair-hair contacts. Finally, the sliding velocity used in the experiments (4 × 10^−6^ m s^−1^) is well below those that can be directly simulated using the current CG-MD framework (1 × 10^−2^ m s^−1^).

**Figure 3:**
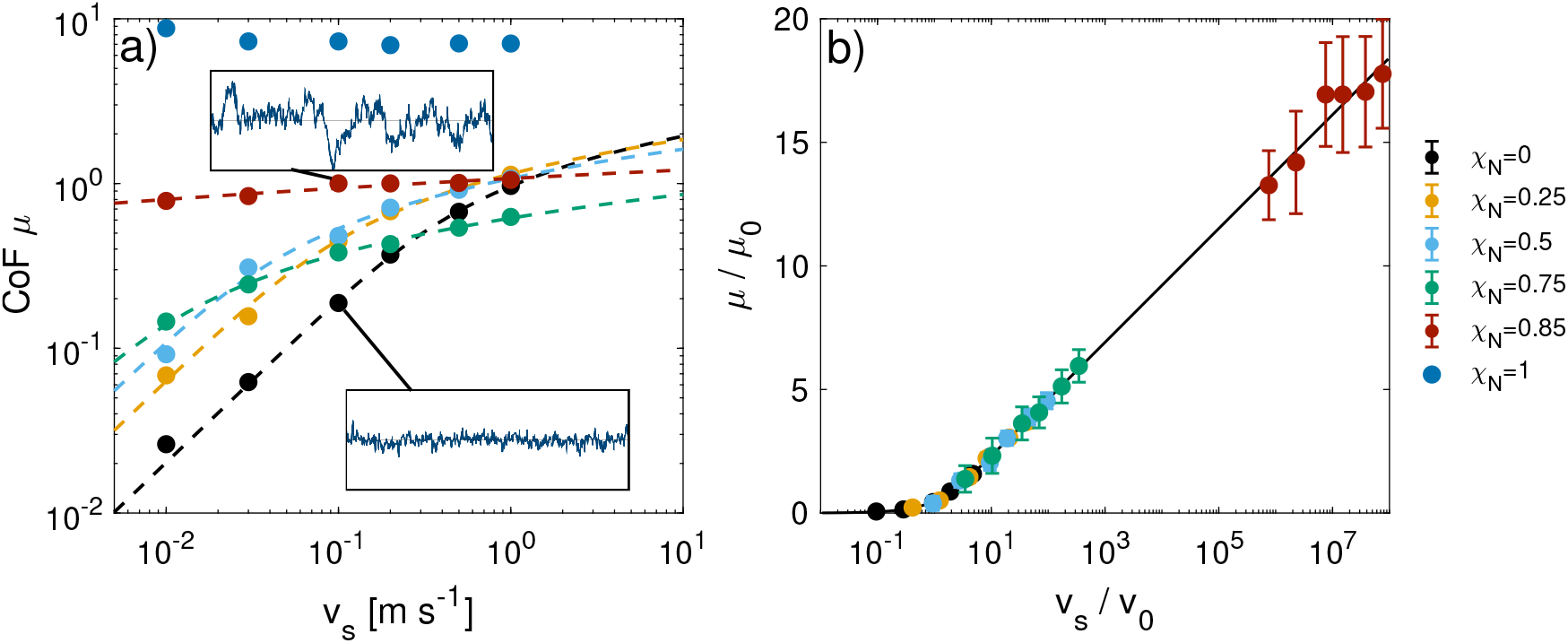
Dry CoF as a function of sliding velocity *v_s_* and monolayer damage at *σ* = 10 MPa in double logarithmic plots. Eq. 1 (dashed lines) is fitted to the data for all damage ratios except χ_N_ = 1.^37^ Uncertainty bars are omitted in a) to improve clarity. b) Shows the same data as a nondimensional representation, collapsed by the fitting parameters *v*_0_ and *μ*_0_. Vertical bars show one standard deviation. Insets in a) show raw friction vs. time signals at the same *x* and *y* axis scale.

For low and moderately damaged hair (χ_N_ < 0.85), the CoF increases sharply with sliding velocity. For higher damage levels, the friction dynamics are much more solid-like, showing only moderate increases in friction force with sliding velocity, as well as stick-slip behaviour. ^91^ The transition from liquid-like, smooth friction towards a more solid-like, stick-slip regime (insets in Fig. 3a and Fig. S4) can be observed by an increase in the standard deviation of the CoF around the mean (shown in Fig. 3b). This is attributed to the reduction in the number of flexible lipids on each surface and increase in the number of inflexible charged groups, which leads to increasingly stiff and heterogeneous surfaces. A crossover is observed at high sliding velocity where the CoF at moderate damage exceeds that at higher damage. For χ_N_ = 0.75, a crossover to the lowest CoF among all degrees of damage at that velocity is observed at *v_s_* > 0.3 m s^−1^. Ultimately bleached (χ_N_ = 1.0) surfaces show an increase in CoF by one order of magnitude compared to the medium bleached case (χ_N_ = 0.85). At this degree of damage, CoF is completely insensitive to sliding velocity, as expected for solid-solid friction. ^91^ The much higher friction for the damaged surfaces is mainly due to the strong electrostatic interactions between the negatively charged sliding surfaces and the positively charged confined counter ions. Similarly high CoFs have also been observed for negatively charged surfaces with strongly confined ionic liquid cations. ^92^

For all systems with χ_N_ ≤ 0.85, the CoF increases logarithmically with sliding velocity, *v_s_*. This behaviour is indicative of a stress-augmented thermal activated (SATA) process,^93^ similar to the Eyring model for viscosity. ^94^ Such behaviour has been observed in many boundary-lubricated contacts protected by adsorbed or grafted monolayers in both experiments^95–97^ and NEMD simulations.^37,38,98,99^ The logarithmic dependency of the CoF on *v_s_* predicted by SATA models generally only holds for intermediate *v_s_*. At very low sliding speeds, a linear relationship is more common, because thermal fluctuations dominate the activation process, as opposed to the external shear stress. ^91,96,97^ An extended SATA model describing both the *v* ∝ ln(*v_s_*) behaviour at high *v_s_* and the *v* ∝ *v_s_* at low has been applied to CoF data from NEMD simulations using both all-atom^37^ and CG^99^ force fields:

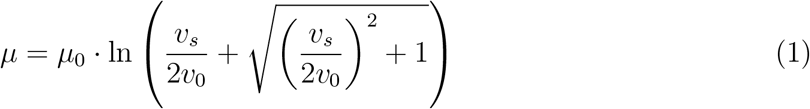

where the reference coefficient of friction, *μ*_0_, and reference sliding velocity, *v*_0_, serve as fitting parameters. The applicability of this model to our data is confirmed for dry hair contacts by means of the dimensionless representation in Fig. 3b). The fitting parameters *μ*_0_ and *v*_0_ are shown in Fig. 4a) and b) as a function of surface damage. The fully damaged case (χ_N_ = 1.0) has been excluded from this fitting since the CoF is insensitive to sliding velocity for this system. The fitting parameters 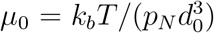 and *v*_0_ = *k*_0_ · *d*_0_ from the SATA model can be related to physically meaningful quantities. Specifically, the distance between barriers, *d*_0_, and the hopping rate constant, *k*_0_, respectfully serve as spatial and temporal indicators to how high and frequent the energy barriers are. The reference CoF and reference velocity both decrease with increasing damage ratio. Fig. 4c)-d) shows that there is an increase in the distance between barriers, *d*_0_, and a decrease in the hopping rate constant, k0, with increasing damage. These changes in *d*_0_ and *k*_0_ indicate that energy hopping events occur more infrequently and with a larger energy barrier distance as the damage ratio increases. This is because the number of lipid islands eventually decreases, while their physical separation increases, as the damage level increases. ^33^ The conformational change to smaller, more widely spaced lipid islands is consistent with the appearance of stick-slip friction at high damage levels (inset in Fig. 3a and Fig. S4).

**Figure 4:**
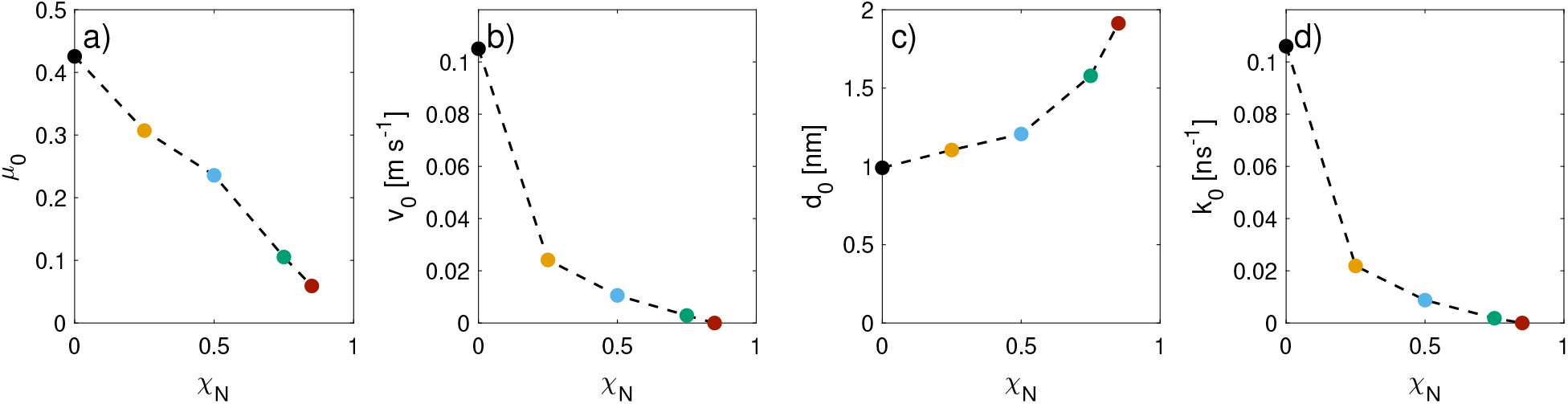
Reference a) coefficient of friction *μ*_0_, b) velocity *v*_0_ and corresponding c) barrier distance *d*_0_ and d) hopping rate constant *k*_0_ from thermal activation model fits^37^ as a function of monolayer damage.

The molecular-scale roughness imposes a physical barrier to overcome during sliding. The *d*_0_ value correlates with the interdigitation between the lipid monolayers on the opposing sliding surfaces (Fig. 2). We quantify the degree of interdigitation by means of the average volume available per bead in the region of direct interdigitation between the two contacts. We find the available volume per bead to increase monotonically from 〈*V_i_*〉 ≈ 0.1 nm^3^ for fully-functionalised monolayers to 〈*V_i_* 〉 ≈ 0.14 nm^3^ for χ_N_ = 0.85 (Fig. S5a). Therefore, more space per CG bead is available at higher damage. This is in contrast to the interdigitation distance, which reaches its peak at intermediate degrees of damage, consistent with a maximum in the expected degree of heterogeneity. ^33^ We also find that the volume per bead (rather than the interdigitation distance) is highly correlated with the barrier distance *d*_0_ from the fits to Eq. 1 (Fig. S5b).

Absolute tilt angles of the vector from the lipid root bead (P_5_) to the terminal bead (C_1_) relative to the xy-plane are shown in the Supplementary Material (Fig. S6a). For the dry contacts, the absolute tilt angle decreases from ≈50°for fully-functionalised surfaces, to ≈25°for medium bleached hair. This suggests that the lipids go from a mostly upright conformation to lying more parallel to the surface as the damage level increases. There is only a slight decrease in the absolute tilt angle with sliding velocity.

We also quantified the orientational tilt angle, which is measured between the projection of the P_5_-C_1_ vector in the xz-plane and the *x* unit vector, as shown in the Supplementary Materials (Fig. S6b). The lipids move from being aligned parallel to the z-direction (which is perpendicular to the hair surface and sliding direction) at low sliding velocity, to being almost completely aligned with the sliding direction at high sliding velocity. Friction is often reported to increase as a function of monolayer alignment in the direction of sliding, which is generally attributed to higher commensurability between the interacting monolayers. ^100,101^ In the current study, however, the monolayers are relatively loosely packed (maximum lipid coverage = 2.7 molecules nm^−2^)^38,102^ and the interface does not form a crystalline, commensurate interface, even when the molecules are aligned with the sliding direction. Indeed, at low damage levels, where high commensurability is most likely due to the highest lipid coverage, there is no stick-slip behaviour evident in the friction versus sliding distance signals (Fig. S4). Molecular alignment seems to have no direct effect on the friction behaviour.

The absence of water in biological systems is rare, even in dry environments. The presence of residual water in the cuticle, ^103^ absorbed water from ambient humidity, ^104^ and the frequent contact of hair with water during washing^11^ demand for the addition of water at the interface, which is discussed in the next section.

### NEMD Simulations of Wet Contacts

#### Squeeze-out Simulations

Squeeze-out simulations are performed at various degrees of chemical hair damage to establish an equilibrium film thickness at the target pressure (10 MPa) for subsequent sliding NEMD simulations. ^38^ The equilibrium number of water units and examples of temporal evolution of the number of polarizable water units (*N_w_*) and contact thickness at *p* =10 MPa are shown in Fig. 5. A steady-state thickness of between 4 and 6 nm is reached after between 13 to 20 ns, depending on the degree of damage. A decrease in the equilibrium thickness and increase in the number of water units is observed with increasing damage. This is because, as the damage level increase, the number of hydrophobic lipids decreases and hydrophilic sulfonate groups become more prevalent on the surface. Stronger intermolecular interactions between the sulfonate groups and the water beads keep more water trapped inside the contact.^105^ The steady state contact thickness decreases until reaching a constant value (*d* = 4.2 nm) at χ_N_ > 0.5 due to the combined effects of a reduced number of lipids and an increase in water content.

**Figure 5:**
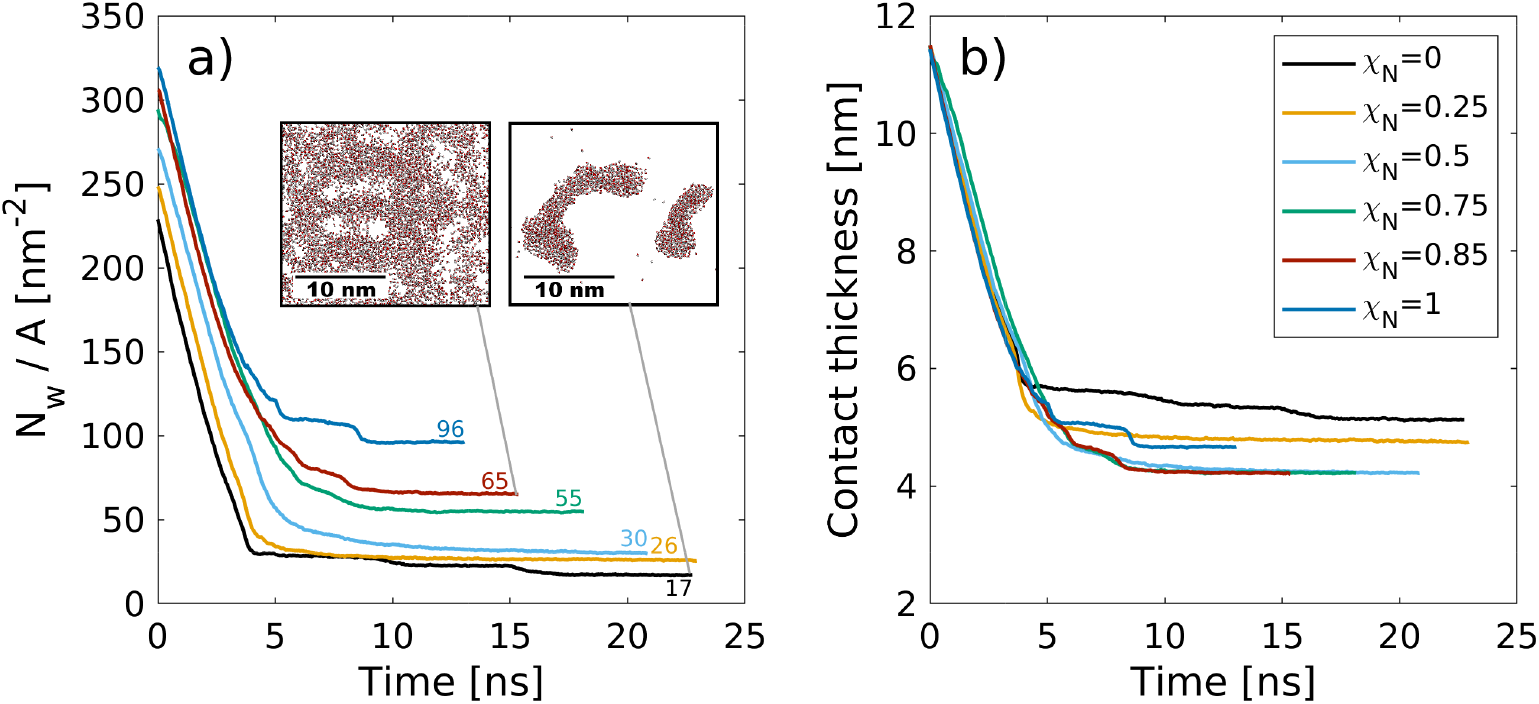
Change in the number of water molecules (4 ·*N_w,CG_*) per surface area *N_w_A*^−1^ a) and the contact thickness between the graphene sheets b) with time during the squeeze-out simulations at a pressure, *σ* = 10 MPa for hair surfaces with different degrees of χ_N_. The molecule number densities per unit area at equilibrium are shown as text labels in a). Insets in a) show snapshots of the confined water beads at the end of the squeeze-out simulations rendered with VMD.^47^ The contact thickness in b) includes the monolayer thickness.

Recent small-angle neutron scattering (SANS) experiments suggest that the cell membrane complex of the hair cuticles in healthy hair contain water clusters with a size of 40 Å.^103^ For formic acid treatment, which removes internal lipids from the hair, the number of such clusters was reduced.^103^ While this transition takes place below the exposed F-layer, the proposed mechanism is comparable to the contacts modeled in this work. In the cell membrane complex, the contact between the 18-MEA monolayers on two neighbouring cuticles is essentially the same as between the outer cuticles of two virgin hairs. In our NEMD simulations, we observe the formation of water clusters of similar size as those observed experimentally^103^ for the fully functionalised systems (χ_N_ = 0). Droplets are energetically favored over thin film formation due to the hydrophobic nature of the two monolayer surfaces. The formation of droplets between hydrophobic monolayers has also been observed in previous NEMD simulations. ^106^ At higher damage levels, water progressively attaches to regions where the lipids have been removed, which is consistent with previous MD simulations^33^ and AFM experiments, ^107^ For the ultimately bleached hair surfaces (χ_N_ = 1.0), an equivalent atomistic water number density per unit area of *N_W_A*^−1^ = 96 nm^−2^ is obtained, which is distributed within six hydration layers normal to the two surfaces. This value is in good agreement with the largest value used in all-atom NEMD simulations that studied the friction of water confined between hydrophilic alkylsilane self-assembled monolayers (*N_W_A*^−1^ = 85 nm^−2^).^108^ For the fully funtionalised surfaces (χ_N_ = 0.0), the final water coverage (*N_W_A*^−1^ = 17 nm’^−2^) is in good agreement with the plateau value (*N_W_A*^−1^ = 20 nm^−2^) obtained from previous atomistic Monte Carlo simulations of water adsorption between hydrophobic graphene surfaces separated by 1.6 nm.^109^

#### NEMD Simulations of Wet Contacts

We used systems containing the equilibrium number of water molecules from the squeeze-out simulations to investigate the friction between the wet hair surfaces with different damage levels. Mass density profiles at *v_s_* = 1 m s^−1^ and *σ* = 10 MPa are shown in Fig. 6 for all χ_N_ ≤ 0.85. The density profiles are again found to be insensitive to the sliding velocity. Strong layering is observed at the interface for both the surface monolayers and water. Such layering is has been found for water in confined contacts in several previous molecular simulation studies.^109,110^ Ordered water layers have also been observed experimentally between adsorbed cetyltrimethyl-ammonium bromide (CTAB) surfactant monolayers and have been shown to facilitate low friction. ^111^

**Figure 6:**
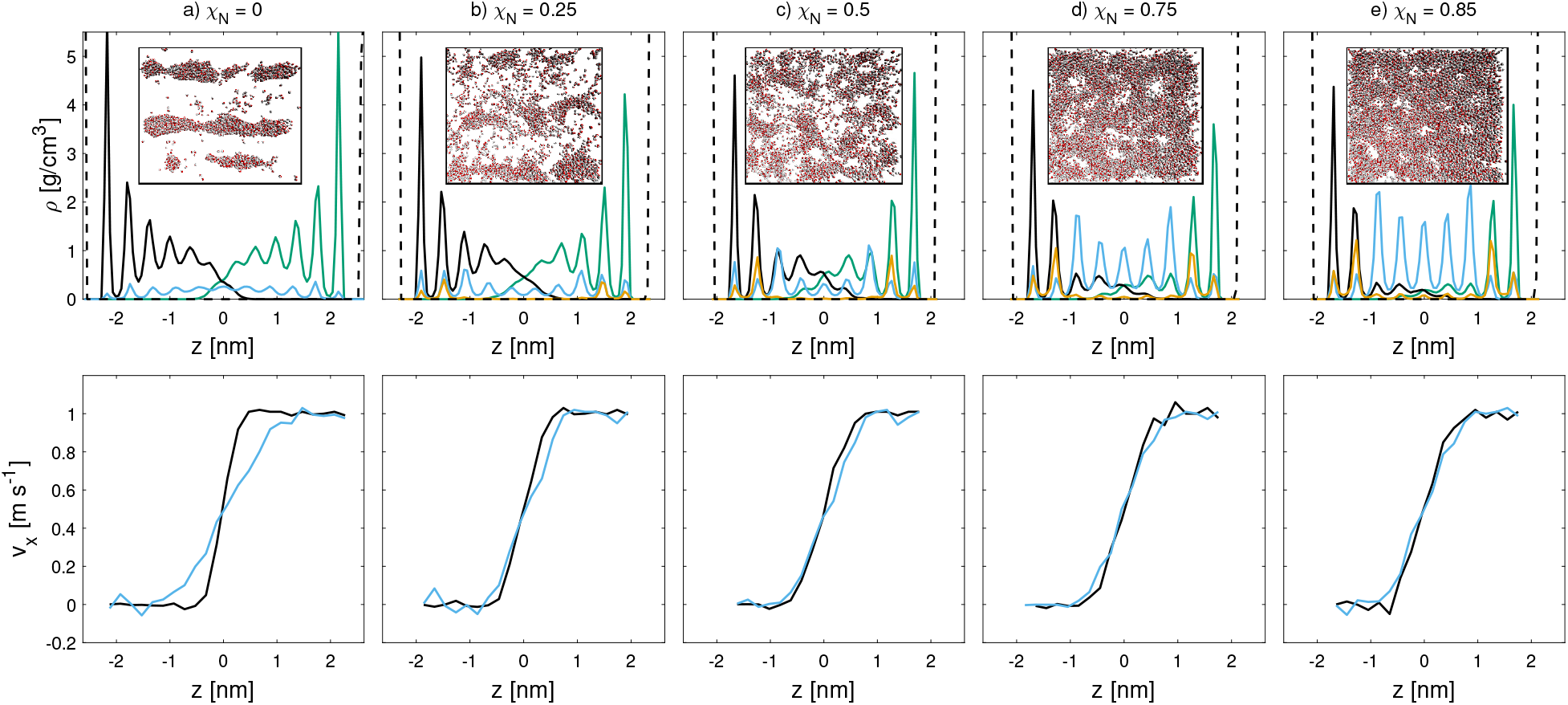
Mass density (top) and velocity (bottom) profiles for wet sliding simulations at *v_s_* = 1 m s^−1^ and *σ* = 10 MPa at different degrees of monolayer damage. Water (blue), lower hair (black), upper hair surface (green), Na^+^ counterions (orange) and base graphene sheets (dashed black) density profiles are shown. Velocity profiles of lipids (black) and water (blue) are shown. Insets show top view snapshots 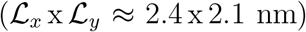 of water inside the contacts. Rendered with VMD. ^47^

The hydrophobic nature of the fully-functionalised hair surfaces (χ_N_ = 0.0) favors the formation of water droplets, which are stretched in the direction of sliding, as evident from the inset in Fig. 6a). Additional snapshots at different sliding velocities are shown in the Supplementary Material (Fig. S7 and Fig. S8). For these undamaged systems, the water droplets penetrate around 2 nm into the monolayers and adhere to the hydrophilic P5 beads at the base of the lipids. At higher χ_N_, thin water films form that are interspersed with lipid islands, as shown by the top view snapshots in Fig. 6. Even at χ_N_ = 0.85, we observe interdigitation between the lipid monolayers on the two opposing surfaces. This is due to swelling of the monolayers, ^33^ which results in mostly upright conformations, even at high damage. At χ_N_ = 1 (not shown), there are no remaining lipids and thus no interdigitation. In this case, there is a complete thin (4 nm) water film between the sulfonate-covered surfaces. Fig. 7 shows the velocity profiles of the lipids among both surfaces and water at the interface. In the fully-functionalised case, there is a slip plane between the lipid monolayers on the opposing surfaces, while the water beads are sheared to the depth that they penetrate the lipid monolayer. This is attributed to the pinning of the water droplets to lower regions of the displaced monolayers. In all other cases, the water beads move at the same velocity as the neighbouring lipid beads.

**Figure 7:**
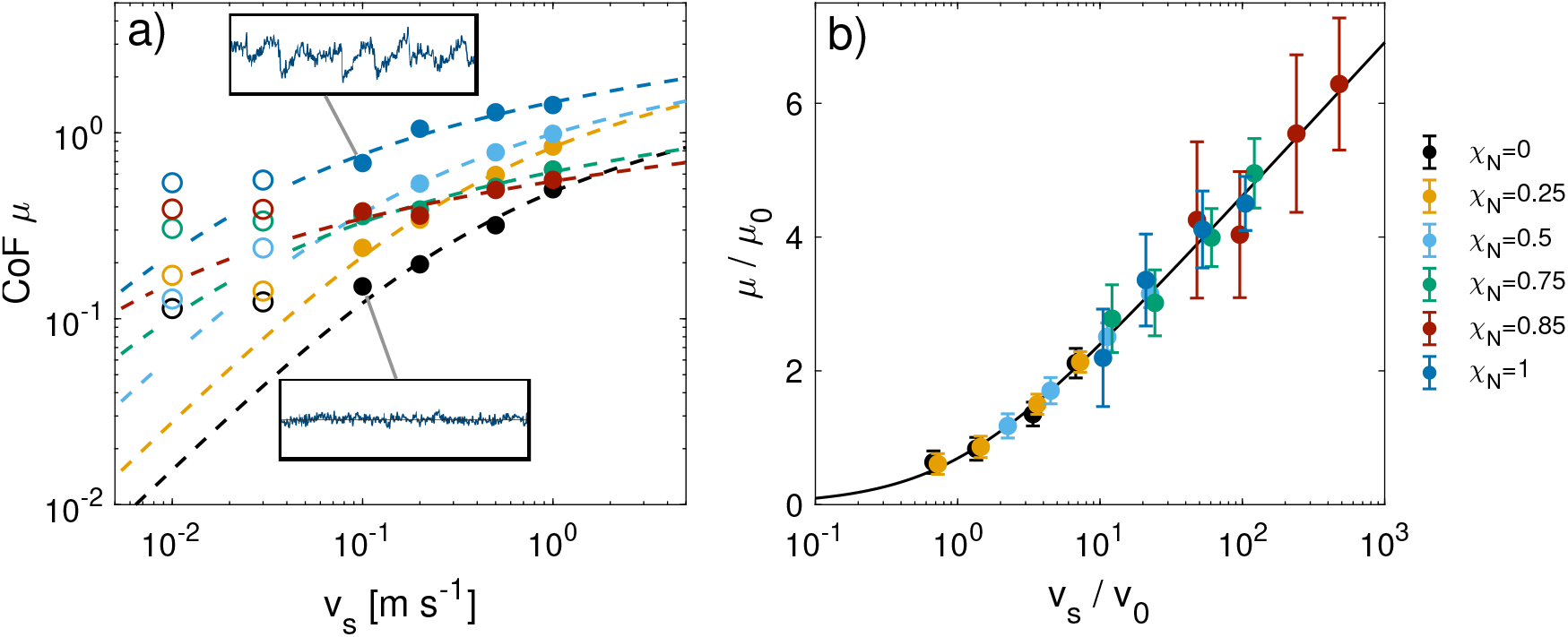
Wet CoF as a function of sliding velocity, *v_s_*, for the different damage ratios at *σ* = 10 MPa in double logarithmic representation. Dashed lines indicate fits Eq. 1.^37^ Open symbols represent data not included in the fits. b) Friction-speed data normalized by the fitting parameters.

For the wet contacts, the absolute tilt angle (Fig. S9a) tends to be invariant to damage at a value of ≈ 50°. This confirms that swelling of the monolayer occurs in water under confinement, with the lipids retaining a more upright conformation at high damage levels. This observation is consistent with our previous single-surface CG-MD simulations. ^33^ As with the dry contacts, there is a slight decrease in the absolute tilt angle at higher sliding velocities. The orientational tilt angle decreases more at lower sliding velocity in the dry systems (Fig. S6b) than the wet systems (Fig. S9b). This suggests that the lipids align with the sliding direction more readily in the dry systems than the wet ones. As with the dry systems, molecular alignment seems to have a negligible effect on the friction behaviour.

The CoF in the wet contacts are shown in Fig. 7a) for the different damage levels as a function of sliding velocity at 10 MPa. At χ_N_ = 0.75 and below, smooth, liquid-like friction signals are observed. Despite the presence of water in the contact, stick-slip friction is observed at χ_N_ = 0.85, as evident from the inset in Fig. 7a). The water film thickness is insufficient to provide complete surface separation, even at the relatively low pressure simulated. Interdigitation is confirmed by the overlapping lipid density profiles in Fig. 6e). In contrast, the fully damaged systems (χ_N_ = 1.0) are fully separated by water distributed among six hydration layers. Nonetheless, stick-slip is observed in this system, as indicated by the standard deviation of the friction force signal of 37% around the mean CoF. This observation is consistent with previous experiments using wet bleached hair-hair contacts. ^30^ The highest CoFs are obtained for the fully damaged system across all sliding velocities. In this case, both surfaces are fully separated by a water film with a thickness of ≈2.6 nm and stick-slip is promoted by surface-bound hydration layers on the highly hydrophilic surfaces. This is consistent with previous NEMD simulations of hydrophilic surfaces. ^112^

At high velocities, a cross-over of μ is observed among different degrees of damage. Apart from the fully damaged surface, the 25-50% damaged systems display the highest friction at high speeds. Here, we need to consider the additional variation of the water content across different χ_N_ as established by the squeeze-out simulations. We believe that this is due to the interplay between two mechanisms as the damage increases: (i) as the surface becomes more hydrophilic, the water content derived from squeeze-out increases, thus providing increased surface separation and reduced lipid interdigitation, which is confirmed by Fig. 6. (ii) At the same time, water penetrates into the lipids as can be seen in Fig. 6, even in undamaged regions, which might increase dissipative contributions.

For *v_s_* > 0.1 m s^−1^, the CoF data is fitted to Eq. 1.^37^ Figure 7b) shows the non-dimensional representation of the CoF included in the fits normalized by the fitting constants *μ*_0_ = *k_B_T*/(*p_N_ ·d*^3^) and *v_0_* =*k*_0_*d*.^37^ The changes in fitting constants as a function of surface damage are shown in the Supplementary Material (Fig. S10). At high *v_s_*, the model accurately describes the observed friction behaviour. At *v_s_* < 0.1 m s^−1^, the CoFs strongly deviate from the fit on the high sliding velocity data. The CoFs become almost independent of the sliding speed below *v_s_* = 0.1 m s^−1^ for all degrees of damage and clearly no longer follow the typical SATA behaviour. The corresponding data points are therefore omitted from the fitting, as indicated by open symbols in Fig. 7a).

One mechanism by which the CoF could exceed that expected from Eq. 1 at low sliding velocities is capillary condensation. Previous AFM experiments using nanoscale silica-silica contacts at moderate relative humidity showed that friction decreased logarithmically with increasing sliding velocity before levelling off. ^114^ Although there is no water-vapour interface in our NEMD simulations, this is a water-lipid interface and increases in adhesion forces have been previously reported for water-oil interfaces. ^115^ In the low-speed regime, nanoscale water droplets (low damage) could give rise to additional cohesive forces through the existence of a well-defined phase boundary. Indeed, previous AFM experiments have shown that water nanodroplets in n-hexadecane increase friction.^116^ Snapshots of these water structures in the fully-functionalised contacts are shown at different sliding velocities in the Supplementary Material (Fig. S7). At full surface damage, no such phase boundary is present in the lateral direction because a thin water film forms.

We quantified the area of contact-bridging water-lipid interface by tracing the perimeter of the water droplet (low damage) or lipid islands (high damage) and multiplying by the average thickness of those structures in the surface-normal direction (Fig. S11). There is a clear transition between intermediate (χ_N_ = 0.5) and high (χ_N_ = 0.75) damage where there is a large increase in the interfacial area. However, the interfacial area weakly increases with sliding velocity and thus cannot be linked to increased capillary adhesion at low speeds, in which case higher a interfacial area would be expected. ^115^ Moreover, capillary condensation would be expected to increase friction on hydrophilic surfaces more than hydrophobic ones, ^34^ which is the opposite to the trend we observe in our NEMD simulations.

Related to capillary effects, freezing of the confined water bridges is another possible explanation for elevated friction at low sliding velocities. Indeed, previous AFM experiments of the phenomenon have observed higher than expected friction at low sliding velocities, as shown in Fig. 7a).^117^ To investigate this effect, the diffusion and structure of the confined water molecules were studied at different sliding velocities and damage levels. The meansquare displacement (MSD) and radial distribution function (RDF) of the central water beads were analysed, as shown in Fig. 8. While the lateral MSD increases linearly with time for highly damaged systems, it approaches a plateau at low degrees of damage, which is indicative of subdiffusion. The more pronounced subdiffusive behaviour at low degrees of damage is partially due to a decrease in confinement lengthscale (Fig. 6), as the majority of the contact is occupied by lipids and water resides within small droplet-like regions. This is in agreement with previous studies of water confined by soft materials,^118^ where the lengthscale of confined water was found to play a role in the tendency of water to show subdiffusion.

**Figure 8:**
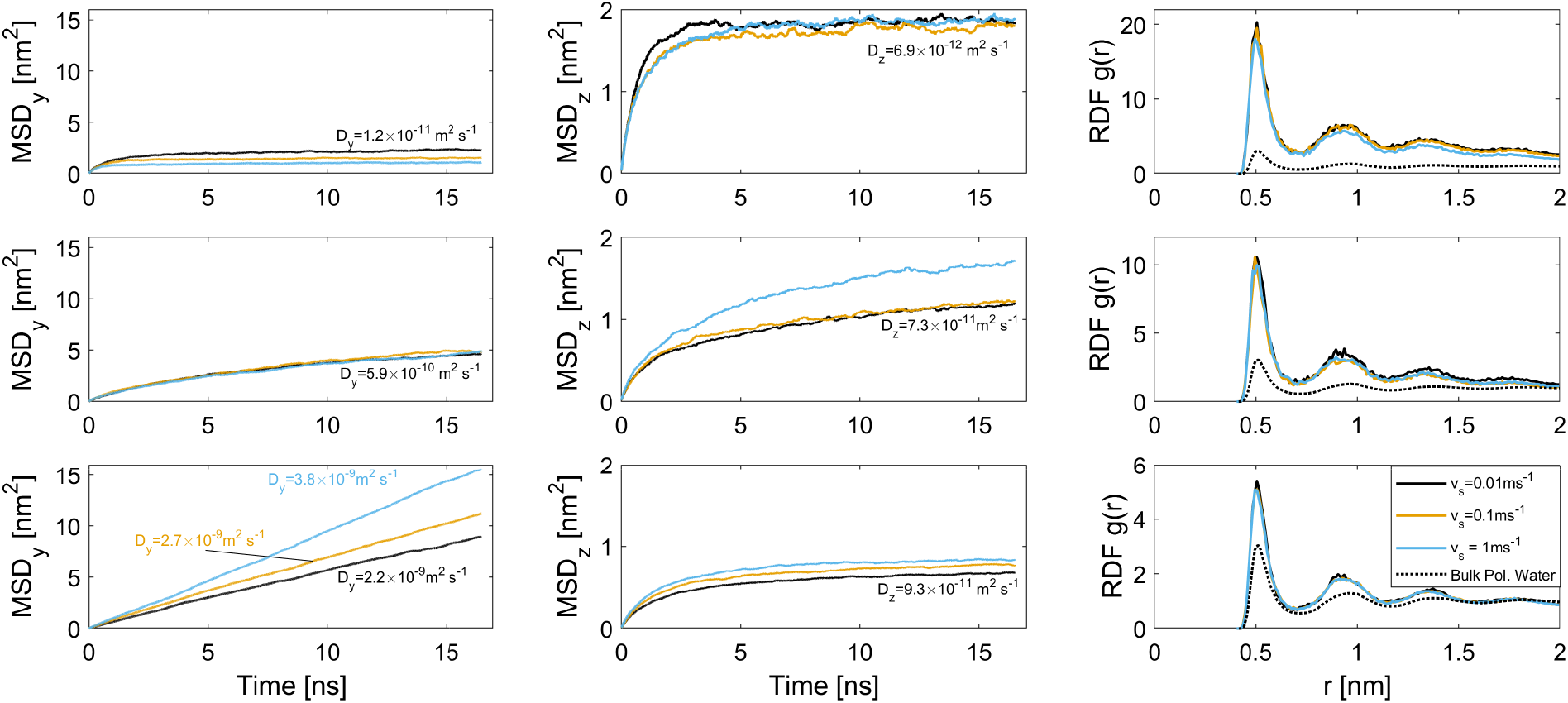
Mean-square displacement (MSD) of the central polarizable water beads perpendicular to the direction of sliding (*y*: left; *z*: middle) at different speeds and for fully-functionalised surfaces (top row), virgin (middle row) and medium bleached hair (bottom row). Here, *t* = 0 ns indicates a time at which the contact has long reached dynamic equilibrium. Approximate diffusion coefficients for subdiffusive configurations are obtained from linear fits for *t* > 10 ns. Right side: Radial distribution function (RDF) of central water beads compared to the RDF in bulk. ^113^

An increase of the lateral MSD in the *y* direction is found with increasing damage. For the medium bleached model hair surfaces (χ_N_ = 0.85), water diffusion coefficients perpendicular to the direction of sliding increase at higher sliding velocity, from *D_y_* = 2.2 × 10^−9^ to 3.8 × 10^−9^ m^2^ s^−1^). These values are similar to those obtained previously for bulk polarizable MARTINI water (*D* = 2.5 × 10^−9^ m^2^ s^−1^)^49^ and liquid water from experiments (*D* = 2.3 × 10^−9^ m^2^ s^−1^).^119^ For the virgin model hair surfaces (χ_N_ = 0.25), long-time lateral diffusion is much slower than for bulk water (*D_y_* = 5.9 × 10^−10^). This reduction is similar to that observed in previous atomistic MD simulations of water inside lipid bilayers. ^120^ For fully-functionalised hair (χ_N_ = 0.0), the lateral diffusion coefficient is more than two orders of magnitude lower (*D_y_* = 1.2 × 10^−11^) than in the bulk. This reduction in lateral diffusion is similar to that observed previously for confined bilayer ice by atomistic MD simulations (*D_y_* ≈ 10^−12^ m^2^ s^−1^).^121^ This implies that the confined water dynamics are solid-like in the fully-functionalised systems. It has also been suggested that surface hydrophobicity might be crucial to confinement-induced liquid-solid water phase transitions^122^ since freezing has not been observed for hydrophilic surfaces in either experiments^123^ or simulations. ^124,125^ Indeed, we note that the lateral diffusion coefficient increases as the damage level, and thus surface hydrophilicity, is increased. In our NEMD simulations, the water mobility is more sensitive to damage than sliding speed and, at low damage, the lateral diffusion coefficients suggest that the water remains in the same state at both low and high sliding velocity.

Structural ordering of the confined water molecules as shown in the RDFs in Fig. 8 is consistent with bulk ordering of polarizable MARTINI water as shown by Sergi et al. ^113^ Although the peak intensities increase somewhat at lower sliding velocity, there are only small shifts in the peak positions for a given degree of surface damage. There do not appear to be additional peaks at larger distances, which would be indicative of long-range order. This suggests that the reduction in water diffusion at low damage levels is due to a confinement induced liquid-amorphous transition, rather than crystallisation. This is consistent with previous atomistic MD simulations of water confined between hydrophobic walls. ^124^ Similar structural investigations were performed for the CG sodium counterions that are adsorbed near the damaged regions of the surfaces, as shown in the Supplementary Materials (Fig. S12). We observe comparable changes in ion diffusion as for water, with a more pronounced mobility in the lateral direction as damage is increased and more diffusion with sliding velocity for medium bleached hair surfaces. Diffusion is generally lower for the counterions than for water, due to the strong electrostatic attraction to the nearby oppositely charged surfaces. The ion RDF is independent of the sliding speed and comparable to the bulk RDF for the cations. ^49^

High friction in damaged hair could arise from strong electrostatic interactions between the negatively-charged sulfonate groups on the surfaces and the strongly-bound, positively-charged sodium counterions confined between them. The counterions are forced to change their distributions from the thermally equilibrated positions by the shear force. This induces an attraction of the charged surface groups and the counterions to pull them back to the original equilibrated positions. As a result, the two repulsive surfaces attract each other via their counterions and show high friction. ^126^

The CoF in wet systems are now compared to the dry counterparts discussed in the previous section. The ratio between the wet and dry contact CoF as a function of sliding velocity at equal pressure is shown in Fig. 9. At sliding speeds of *v_s_* = 0.1 m s^−1^ and above, wet contacts render a friction reduction for all of the systems. The largest reduction at all sliding velocities is for the ultimately bleached (χ_N_ = 1.0) system. At low sliding velocities, wet friction forces exceed dry friction forces for the systems with low damage levels (χ_N_ < 0.85).

**Figure 9:**
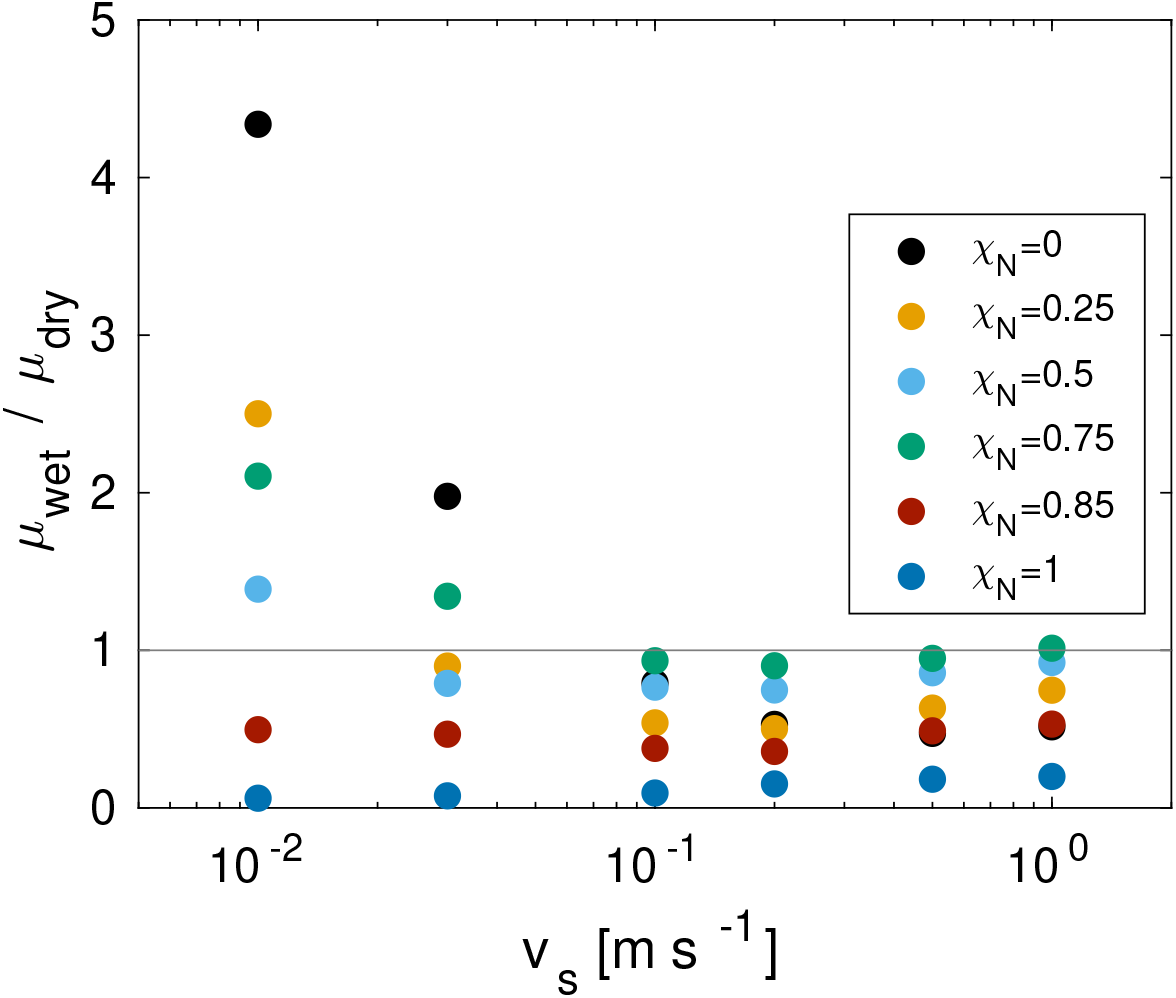
Ratio between wet and dry CoF as a function of sliding speed *v_s_* and monolayer damage χ_N_ at *σ* = 10 MPa.

This is attributed to increased adhesive contributions of solid-like interfacial water droplets confined between the surface-bound lipids. Conversely, large friction reductions are found for the fully damaged surfaces because of the thin water film, which provides sufficient separation to lubricate the contact interface.

Previous experiments have also shown higher friction for wet hair than dry hair. ^8^ Previously, this has been attributed to microscale effects, specifically the swelling of the cuticles leading to increased contact area and rougher surfaces. ^11^ Such mechanisms are not captured by our CG-MD model, apart from moderate swelling of the monolayer.^33^ Our NEMD simulations suggest that solidification of confined water droplets^117^ is an additional nanoscale mechanism that could lead to increased adhesion and friction between wet hairs.

#### Comparison between NEMD Simulations and AFM Experiments of Wet Contacts

The friction behaviour from the wet NEMD simulations is compared to AFM experiments on real hair in humid air^13^ and new CCP AFM experiments with the biomimetic surfaces in water. The shear and normal forces for virgin and damaged hair obtained in the previous AFM experiments^13^ were converted to stresses *τ_0_* using JKR theory^86^ with previously measured surface energies for hair. ^127^ The CCP AFM shear stress and normal stress values are converted from the measured forces using the extension to JKR theory^86^ for thin coatings due to Reedy. ^87^ The experimentally-measured normal and friction forces, prior to conversion to stresses, are shown in the Supplementary Materials (Fig. S13).

In the NEMD simulations (Fig. 7), a monotonic increase of the CoF with the damage ratio is observed at the lowest sliding velocity (*v_s_* = 1 × 10^−2^ m s^−1^). For the fully-functionalised surfaces (χ_N_ = 0.0) at 10 MPa, we obtain a CoF (σ/τ) of *μ*=0.11±0.04, which increases to a maximum of *μ*=0.54±0.21 in the case of fully damaged surfaces (χ_N_ = 1.0). The biomimetic surfaces using CCP AFM functionalised with C_18_-C_18_(fully-functionalised) and SO_3_^−^-SO_3_^−^ (ultimately bleached) yield CoFs of *μ*=0.023±0.002 and *μ*=0.67±0.03, respectively. This means that the CoF of the ultimately bleached contact is ~ 29 times greater than the fully-functionalised contact. Therefore for surfaces representative of ultimately bleached hair, i.e. with all of the lipids removed, the agreement in the CoF from the experiments and NEMD simulations at a single pressure is excellent (−19 %). For the fully-functionalised surfaces, the CoF is around five times larger higher in the simulations than the experiments. This appreciable difference could potentially be explained by a non-negligible load-independent adhesive contribution to the friction force, known as the Derjaguin offset. ^90,102^ To quantify this contribution, we performed additional squeeze-out and NEMD simulations at elevated pressures (*σ* = 20 – 50 MPa). In these simulations, a sliding velocity of *v_s_* = 0.1 m s^−1^ was applied.

Fig. 10 shows the shear stress, *τ*, as as a function of normal stress, *σ*, for (a) the NEMD simulations and (b) CCP AFM experiments. In both the experiments and NEMD simulations, there is a linear increase in *τ* with *σ*, as expected from Amontons’ friction law. ^90^ However, in some cases, there is a non-negligible intercept, which is indicative of an adhesive contribution that can be quantified though the Derjaguin offset.^90,102^ The CoFs (gradient) and Derjaguin offsets (intercept) shown in Table 1 are extracted from the linear fits shown in Fig. 10 to the equation: *τ* = (*σ* · *μ*) + *τ*_0_, where *τ*_0_ is the Derjaguin offset.^90,102^

**Figure 10:**
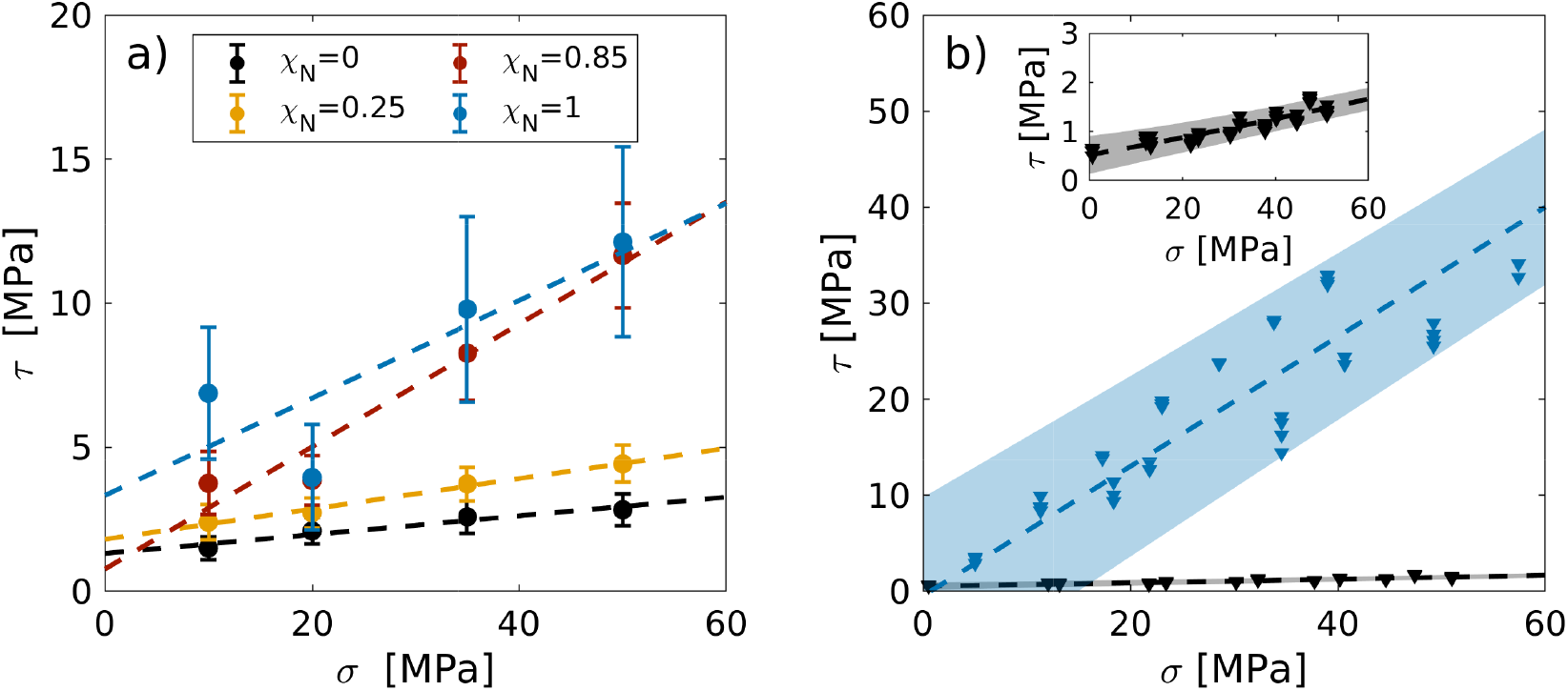
Shear stress, *τ*, as a function of normal pressure, *σ*, from a) wet NEMD simulations and b) biomimetic CCP AFM experiments in water. Experiments for C_18_-C_18_ (χ_N_ = 0) and SO_3_^−^-SO_3_^−^ (χ_N_ = 1) surfaces. Vertical bars for the NEMD data represent the standard deviation of the friction signals. The inset in b) shows the magnified C_18_-C_18_ friction data. Linear fits for NEMD and experimental data are shown as dashed lines. Corresponding prediction intervals on the experimental data with 95% confidence are shown as shaded areas. Experimental pressures are obtained from the extension to JKR theory^86^ for thin coatings due to Reedy. ^87^

**Table 1:** CoF, *μ*, and Derjaguin offset, *τ*_0_, results from wet NEMD simulations (*σ* = 10 – 50 MPa) in fully-functionalised (χ_N_ = 0.0), virgin hair (χ_N_ = 0.25), damaged hair (χ_N_ = 0.85) and ultimately bleached hair (χ_N_ = 1.0) contacts. Experimental results from CCP AFM using C_18_-C_18_ (fully-functionalised) or SO_3_^−^-SO_3_^−^ (ultimately bleached) contacts (*σ* =1 – 60 MPa) and previous AFM results for silicon nitride tips on virgin and damaged hairs (*σ* = 6 – 47 MPa) in humid air (RH ≈ 50%).^13^

| | Exp. $\mu$ | Sim. $\mu$ | Exp. $\tau_0$ [MPa] | Sim. $\tau_0$ [MPa] |
| --- | --- | --- | --- | --- |
| Fully-functionalised (Wet) | $0.023 \pm 0.002$ | $0.033 \pm 0.027$ | $0.52 \pm 0.1$ | $1.3 \pm 0.9$ |
| Ultimately bleached (Wet) | $0.67 \pm 0.03$ | $0.169 \pm 0.149$ | $-0.5 \pm 1.8$ | $3.3 \pm 11.0$ |
| Virgin hair (Air)<sup>13</sup> | $0.03 \pm 0.01$ | $0.053 \pm 0.017$ | 2.3 | $1.8 \pm 0.6$ |
| Damaged hair (Air)<sup>13</sup> | $0.13 \pm 0.05$ | $0.211 \pm 0.147$ | 5.7 | $0.8 \pm 4.8$ |

The uncertainty of the Derjaguin offsets from our NEMD simulations is high, particularly for the damaged hair and ultimately bleached hair contacts. In the CCP AFM experiments, the Derjaguin offset is negligible for the ultimately bleached system and small (*τ*_0_ = 0.52 ±0.1 MPa) for the fully-functionalised system, suggesting some adhesion. The NEMD simulations also give a non-negligible Derjaguin offset for the fully-functionalised case (*τ*_0_ = 1.3 ± 0.9 MPa). A certain degree of adhesion for the fully-functionalised system in water is expected due to the hydrophobic nature of the lipid monolayers. ^128^

The Derjaguin offsets in the previous AFM experiments with hair are somewhat larger than those found in our NEMD simulations and CCP AFM experiments. ^13^ The Derjaguin offset was observed to be larger for damaged hair than virgin hair. ^13^ This is the opposite trend to that observed in the CCP AFM experiments with the biomimetic surfaces.

The trends between the different damage levels in the CoF values from the NEMD simulations are in qualitative agreement with the CCP AFM experiments, as shown in Fig. 10. For the fully functionalised surfaces, the CoF is overestimated (+43 %), while for the ultimately bleached surfaces, the CoF (−75 %) is underestimated by the simulations compared to the experiments. In previous NEMD simulations with *MARTINI*, the intermonolayer friction of lipid bilayers was underestimated by around an order of magnitude compared to experiment. ^43^ Thus, the level of agreement with experimental friction results achieved here with a coarse-grained force field can be considered acceptable.

The CoF of the fully-functionalised hair mimic surfaces in CCP AFM (*μ*=0.023±0.002) is very similar to that obtained with a silicon nitride AFM tip on hair (*μ*=0.03±0.01).^13^ However, the CoF for the ultimately bleached hair mimic (*μ*=0.67±0.03) was much higher than that between a silicon nitride AFM tip and damaged hair (*μ*=0.13±0.05).^13^ This is probably due to the fact that only the substrate was charged in the previous AFM experiments, rather than both as in the current CCP AFM experiments, leading to weaker counterion trapping. ^126^

For the single-pressure NEMD simulations of the intermediate systems, which are representative of virgin (χ_N_ = 0.25) and medium bleached (χ_N_ = 0.85) hair contacts (Fig. 7), CoFs of *μ* = 0.14±0.05 and *μ* = 0.39±0.12 were respectively obtained at 10 MPa and the lowest sliding velocity (*v_s_* = 1 × 10^−2^ m s^−1^). Therefore, the CoF for medium bleached hair is more than twice that of virgin hair. For the variable-pressure (10-50 MPa) NEMD simulations (Fig. 10), the COFs were *μ*=0.053±0.017 for virgin hair and *μ*=0.211±0.147 for medium bleached hair. Thus, the CoFs for both systems are somewhat overestimated in the NEMD simulations compared to the previous AFM experiments (+50 %).^13^ However, the trends between the different damage levels are in excellent agreement with the experiments, with the CoF for damaged hair being approximately four times higher than virgin hair (Table 1). ^13^ This prediction is also consistent with several other previous AFM experiments in humid air and water environments for both bleached and KOH-damaged hair. ^10,22^

Overall, there is good agreement between the friction results from the nanoscale AFM on hair,^13^ the CCP AFM experiments with biomimetic surfaces, and the NEMD simulations (Table 1). The CoF is generally higher in the simulations than the experiments (other than the ultimately bleached system). This is probably due to higher sliding velocity necessary in the NEMD simulations (*v_s_* = 1 × 10^−2^ m s’^1^) than the hair mimic CCP AFM (*v_s_* = 2 × 10^−7^ m s^−1^) and hair-tip AFM (*v_s_* = 2 × 10^−6^ m s^−1^) experiments.^13^ The agreement of the NEMD data with experiments suggests that nanoscale friction increases associated with hair damage are primarily induced by the changes in surface chemistry rather than changes in microscale surface roughness, as suggested from previous experimental results. ^13^ Indeed, it has recently been shown using an AFM relocation method that the nanoscale roughness of hair does indeed not change significantly for a single 10-minute bleaching treatment.^129^

## Conclusion

We have investigated the nanoscale friction between model hair surfaces at different degrees of damage using coarse-grained NEMD simulations and CCP AFM experiments. Biomimetic surfaces were produced by functionalising the silica surfaces with either octadecyltrimethoxysilane, to represent the outer 18-MEA monolayer on virgin hair, or sulfonate groups, to represent the oxidised cysteic acid groups on ultimately bleached hair surfaces. In the CCP AFM experiments in water, we observed much higher CoFs for the ultimately bleached hair model surfaces than the virgin hair model surfaces. This observation is in agreement with previous AFM experiments of silicon nitride tips sliding on virgin and damaged hair.

In the NEMD simulations, we measured the friction between surfaces representative of virgin, medium bleached, and ultimately bleached hair, as well as intermediate degrees of damage. Both dry and wet contacts were considered over a wide range of sliding velocities. For dry and wet contacts at high sliding velocities, we find a near-logarithmic dependency of friction on sliding velocity, which is indicative of a SATA process. We successfully applied an Eyring-like extended SATA model to our NEMD simulation data. For wet contacts, we observe a departure from the SATA behaviour at low sliding velocities as the CoF levels off and approaches a constant value. Friction reductions due to the presence of water in the contact is observed for medium bleached hair at all applied sliding velocities, but only at high sliding velocity for virgin hair. For the hydrophobic virgin hair models, the confined water molecules appear to become more solid-like at low sliding velocities, which could explain the increase in friction relative to the dry systems. At low sliding speeds, we observe a monotonic increase of friction forces with increasing chemical damage. This is in good agreement with AFM experiments on both real hair surfaces and the CCP AFM measurements with the biomimetic surfaces. We also performed additional NEMD simulations at varied pressure to quantify the zero-pressure adhesive contribution to the shear stress, known as the Derjaguin offset. In both the NEMD simulations and CCP AFM experiments, a non-negligible Derjaguin offset was observed, due to hydrophobic. Moreover, the fourfold increase in CoF observed in the varied pressure NEMD simulations moving from virgin to damaged hair is consistent with previous AFM experiments. This observation demonstrates that friction increases of chemically damaged hair at the nanoscale are controlled by the modified surface chemistry, rather than roughness changes or subsurface damage. While this is not a mutually exclusive relationship, we find friction increases in total absence of any roughness effects above the molecular scale. We expect the experimental and computational model surfaces proposed here to be useful for the screening of the tribological performance of hair care formulations.

## Data accessibility

Additional figures are provided in the Supplementary Material, as referenced in the main text. NEMD input files and initial configurations for LAMMPS have been deposited on a dedicated repository, available at https://github.com/erikweiand/cg-hair-nemd.

## Authors’ contributions

E.W. methodology, investigation, data curation, formal analysis, visualization, writing – original draft; J.P.E. conceptualization, methodology, supervision, writing – original draft; Y.R. conceptualization, methodology, investigation, data curation, formal analysis, writing – review & editing; P.H.K. conceptualization, project administration, writing – review & editing; S.H.P. methodology, writing – review & editing; F.R.R. project administration, writing – review & editing; S.A.U. conceptualization, supervision, writing – review & editing; D. D. conceptualization, funding acquisition, resources, supervision, project administration, writing – review & editing.

## Acknowledgements

We thank Sumanth Jamadagni and Andrei Bureiko (Procter and Gamble) for useful discussions.

