## Supplementary Material for "Nanoscale Friction of Biomimetic Hair Surfaces" (from another version of this preprint: version 1, 2022-09-30)

### 1 Contact pressure estimates

#### 1.1 AFM using hair

A crossed-cylinder contact is assumed for a contact between two hair fibres. The contact radius $a$ from Hertzian theory in this case is given by:

$$a^3 = \frac{3RF_N}{4E},$$

where $R$ is the fibre radius, $F_N$ is the applied normal load and $E$ is the Young's modulus of a fibre. Here, a Young's modulus of $E = 0.9$ GPa or $E = 0.5$ GPa is used, which is in accordance with deformation experiments of soaked, virgin and chemically damaged hair, respectively.[1] With an estimate of the hair diameter $d = 2R = 75\mu$m, the remaining variable is the normal load. Using a value of $F_N = 0.1$ mN, which is similar to previous AFM experiments of hair-hair friction,[2,3] we obtain a contact pressure estimate of $p_N = 10$ MPa.

This is a relatively low load compared to hair manipulation scenarios such as rubbing with a finger ($F_N \approx 50 - 100$ mN),[4] but is deemed suitable as a baseline to describe friction between two individual hair fibres in AFM experiments. Estimated pressures as a function of normal load for virgin and damaged hair are shown below.

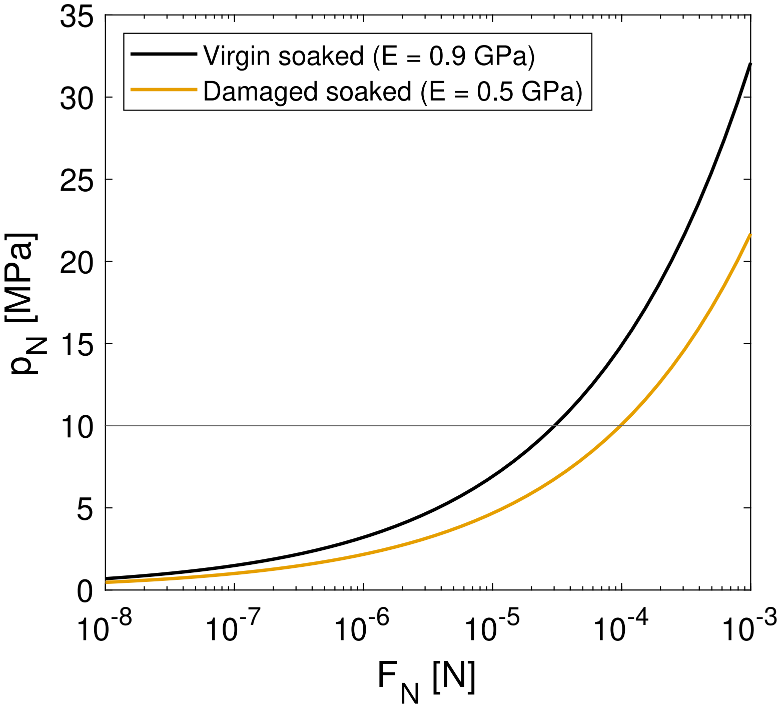


**Figure S1** Pressure *versus* load curves for virgin and damaged hair-on-hair contacts obtained from Hertz theory for a range of normal loads from 10 nN to 1 mN. Horizontal line represents the pressure chosen for the NEMD simulations ($p_N = 10$ MPa).

#### 1.2 AFM using biomimetic surfaces

We also estimated the Hertzian pressure in the chemical colloidal probe (CCP) AFM experiments using the $C_{18}$-functionalised biomimetic surfaces. We used a Young's modulus of $E = 73$ GPa and Poisson ratio of $\nu = 0.25$ for both the silica probe and oxidised silicon substrate,[5] obtaining pressures of $p_{\rm H} < 347$ MPa over the applied load range ($F_N < 3$ $\mu$N), as shown in Fig. S2a).

In addition to Hertz theory, we compared three other methods to estimate the contact pressure in the CCP AFM experiments using the biomimetic surfaces that account for adhesion and/or deformation of the monolayer coatings. Johnson-Kendall-Roberts theory (JKR)[6] was used to account for adhesion between the monolayers ($w_0 = 26.4$ mJ m$^{-2}$). This is similar to the surface free energy measured for 1-hexadecanethiol monolayers on silicon wafers in water.[7] Using the same mechanical properties for the silica surfaces as above, the pressures decrease to $p_{\mathrm{JKR}} < 200$ MPa. This pressure range is comparable to previous JKR estimates from CCP AFM experiments between a non-functionalised silica glass sphere and flat silica surface.[8]

To account for the deformation of the compliant monolayer, we also developed a Newton solver with respect to the contact radius to optimize the constitutive Equation (A15) due to Perriot and Barthel.[9] The contact is idealized as a single coating indented by a rigid spherical probe, therefore neglecting adhesion. A Poisson ratio $\nu_{\mathrm{coating}} = 0.4$ and Young's modulus $E_{\mathrm{coating}} = 0.2$ GPa were used for the coating. This is similar to the Young's modulus measured using AFM for stearic acid monolayers prepared using the Langmuir-Blodgett method on silicon wafers.[10] The resulting pressure range is $p_{\mathrm{PB}} < 146$ MPa for the octadecyltrimethoxysilane functionalized surfaces.

Finally, we used the method proposed by Reedy to account for both deformation of the monolayer and adhesion.[11] Using the same parameters as above, we obtain pressures of $p_{\mathrm{R}} < 79$ MPa. A comparison of the pressure *versus* load curves obtained using the Hertz, JKR, Perriot-Barthel, and Reedy methods is shown below. The same formalism was repeated for the sulfonic acid-functionalised biomimetic surfaces, as evident in Fig. S2b). Hertzian pressures do not change compared to the $C_{18}$-$C_{18}$ contacts as the underlying substrate is the same.

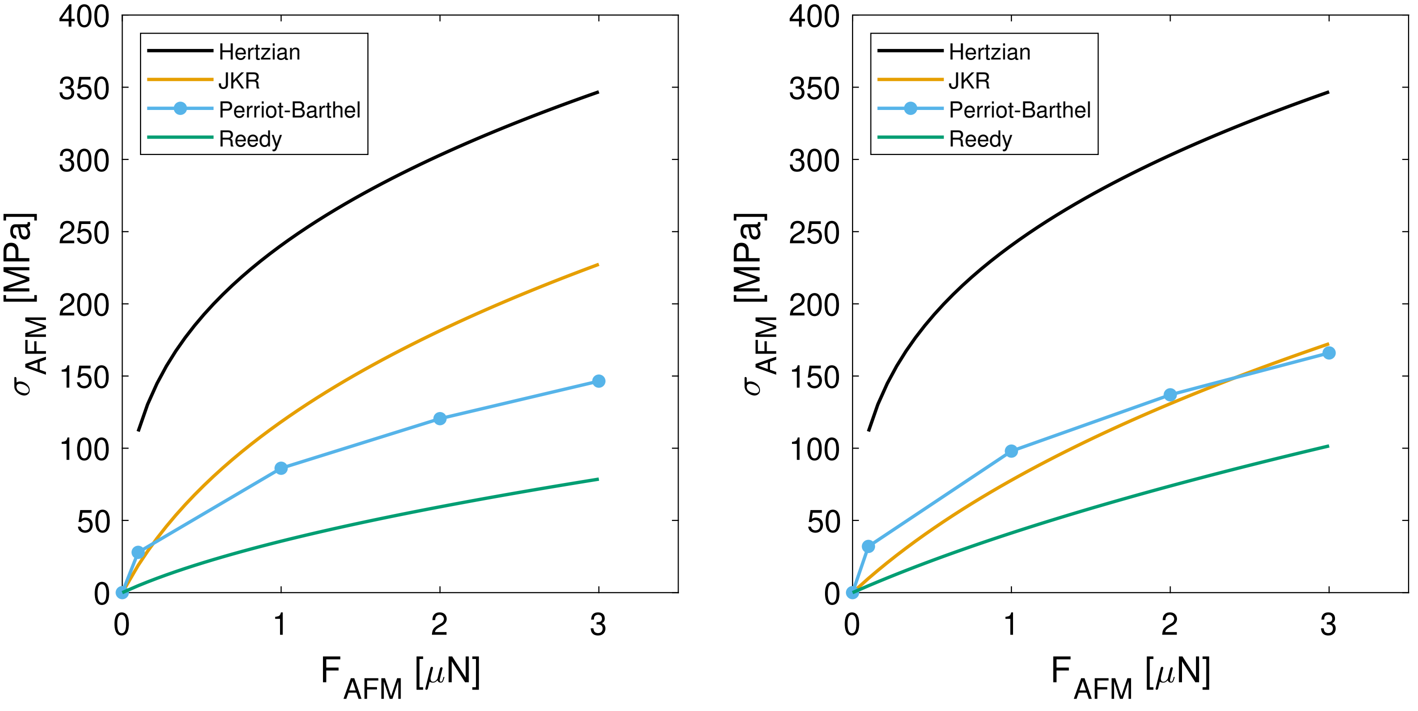


**Figure S2** Pressure *versus* load curves for biomimetic CCP AFM using Hertz theory, JKR theory[6] and the procedure presented by Perriot-Barthel.[9] for a) $C_{18}$-$C_{18}$ surfaces and a) $SO_3^-$-$SO_3^-$ surfaces. Adhesion forces between the monolayers are used for the JKR estimate. Coating deformation properties of the monolayers are applied for the Perriot-Barthel estimate. Adhesion and monolayer deformation are both considered in the Reedy method.

#### 2 MARTINI water viscosity

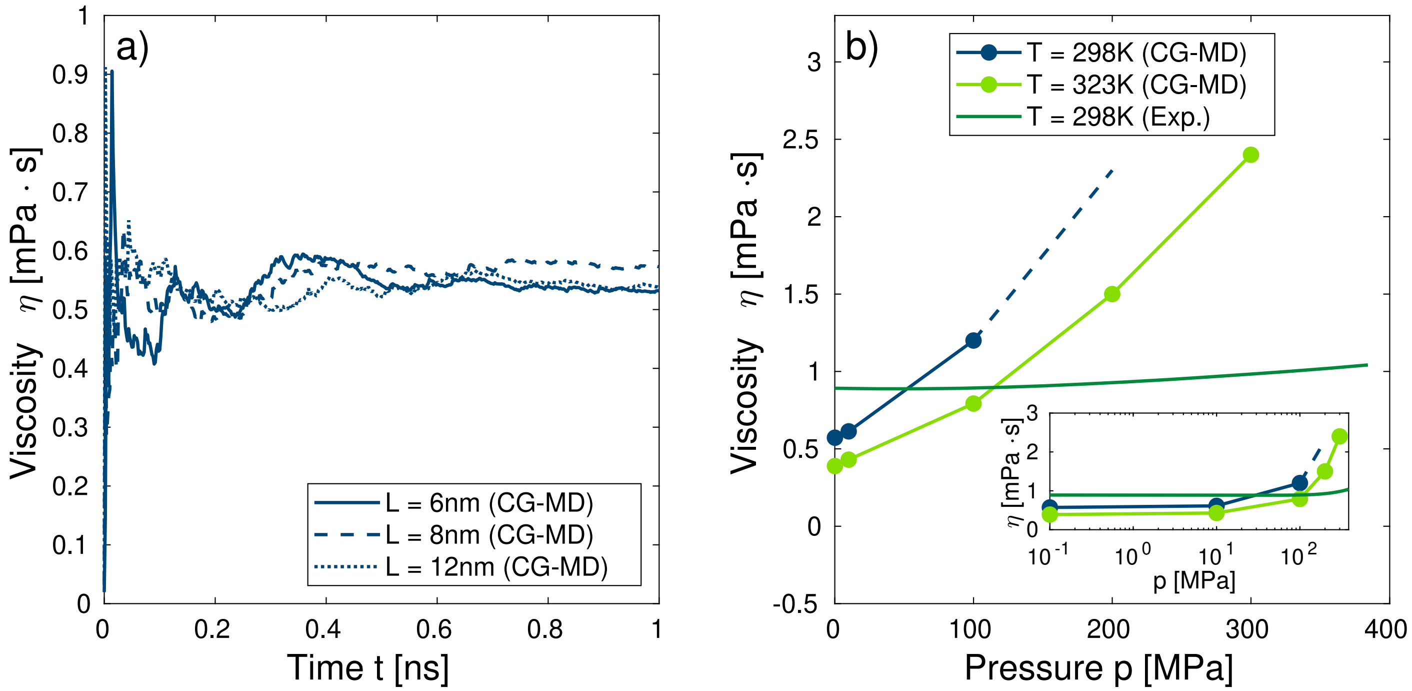


**Figure S3** Dynamic viscosity of bulk system of polarizable MARTINI water beads[12] a) at $p = 1$ atm and $T = 300$ K and three different simulation box sizes ($\mathscr{L} = 6, 8, 12$ nm) and b) at different pressures with constant box size ($\mathscr{L} = 8$ nm). The Green-Kubo formalism[13,14] has been applied to obtain the bulk viscosities. Experimental reference data at $T = 298$ K[15] are shown for comparison.

### 3 Dry/wet friction signals

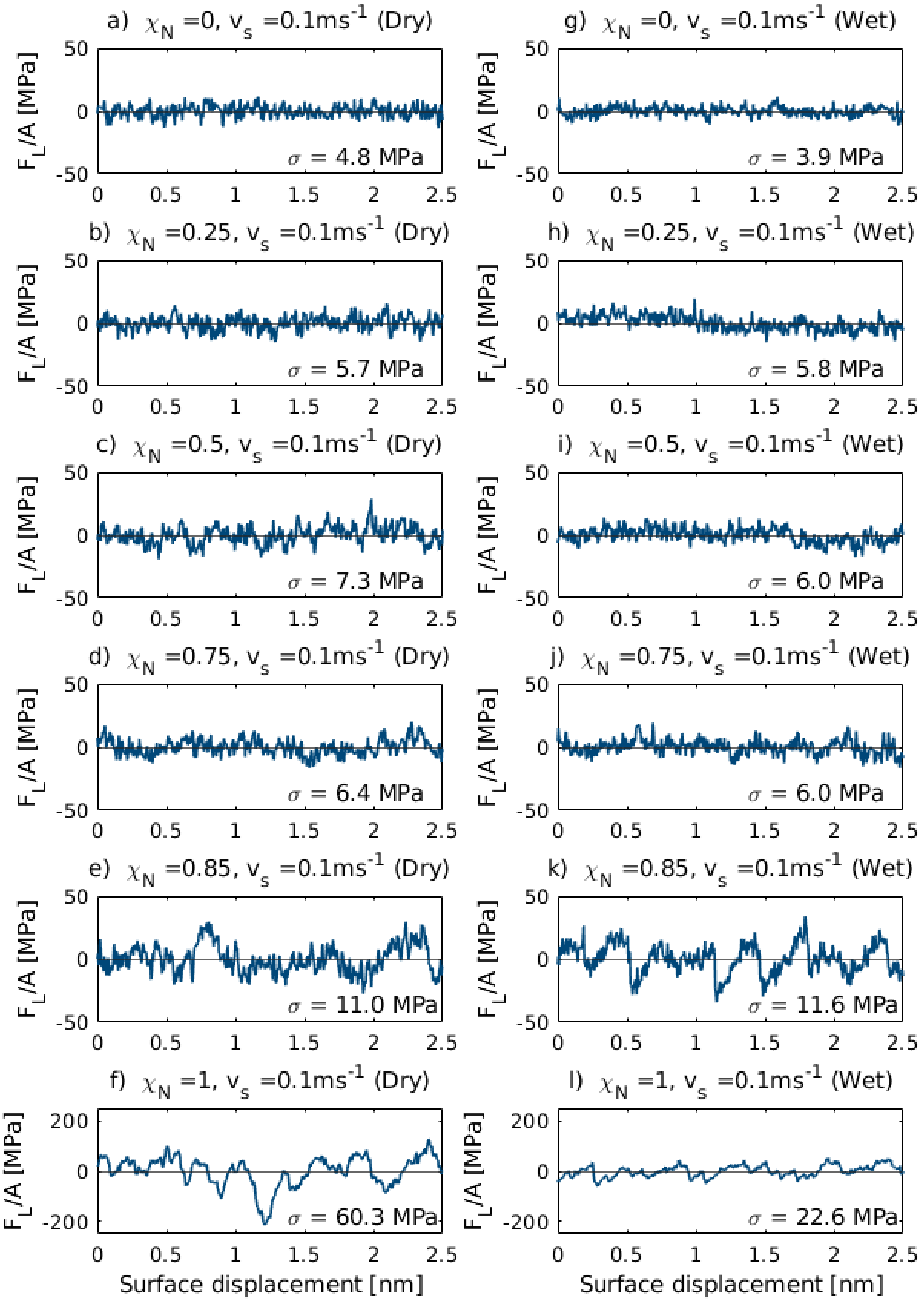


**Figure S4** Mean-removed friction signals, $F_L/A - \langle F_L/A \rangle$, at $v_s = 0.1$ m s$^{-1}$ for dry (left column) and wet (right column) systems. Note that the fully damaged friction signals in f) and l) have increased $y$ axis limits. The standard deviation, $\sigma$, is given for each signal.

### 4 Dry interdigitation volume and SATA energy barrier

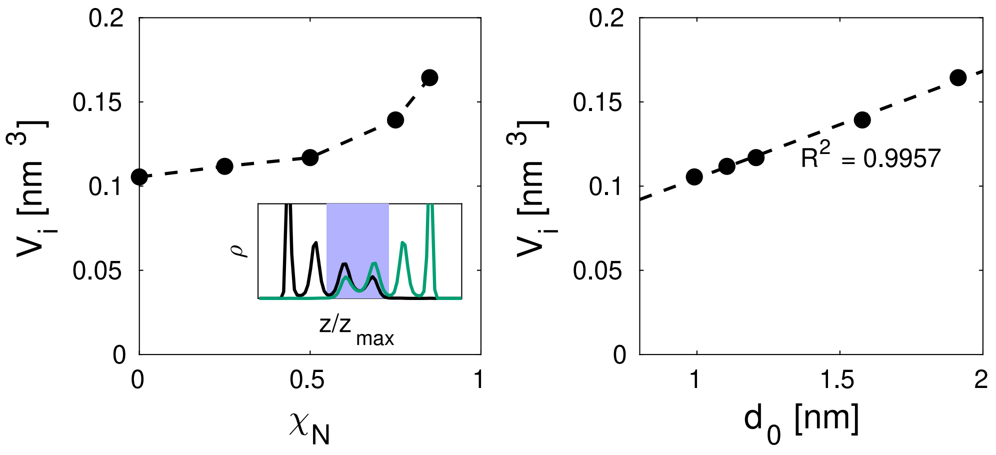


**Figure S5** Average volume per CG bead in the region of interdigitation as a function of a) damage and b) barrier distance $d_0$. An example of the region of interdigitation is highlighted in the inset in a). A linear fit (dashed line in b) confirms that interdigitation directly correlates with the theoretical predictions of a physical barrier to sliding.

### 5 Dry lipid tilt angles

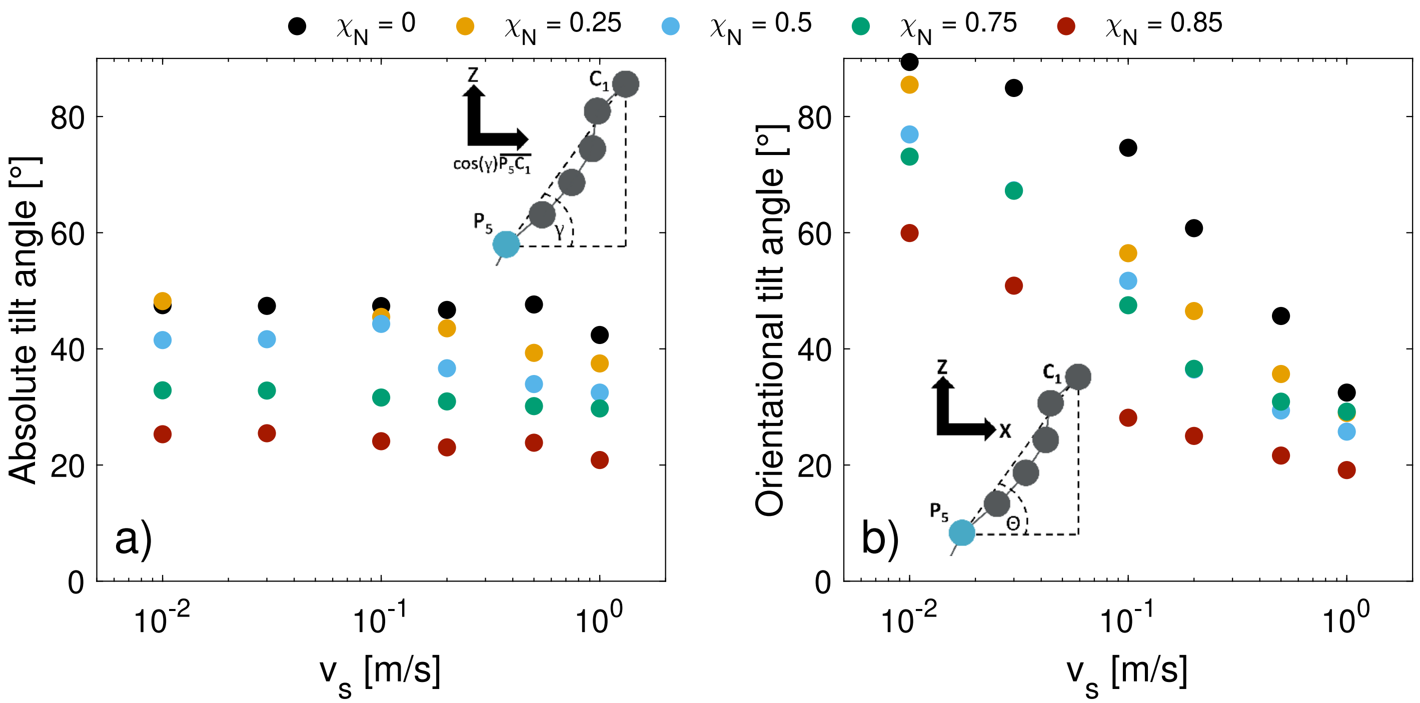


**Figure S6** Absolute lipid tilt angles and orientational tilt angles relative to the direction of sliding for dry contacts as a function of sliding speed and surface damage.

### 6 Water droplet structures

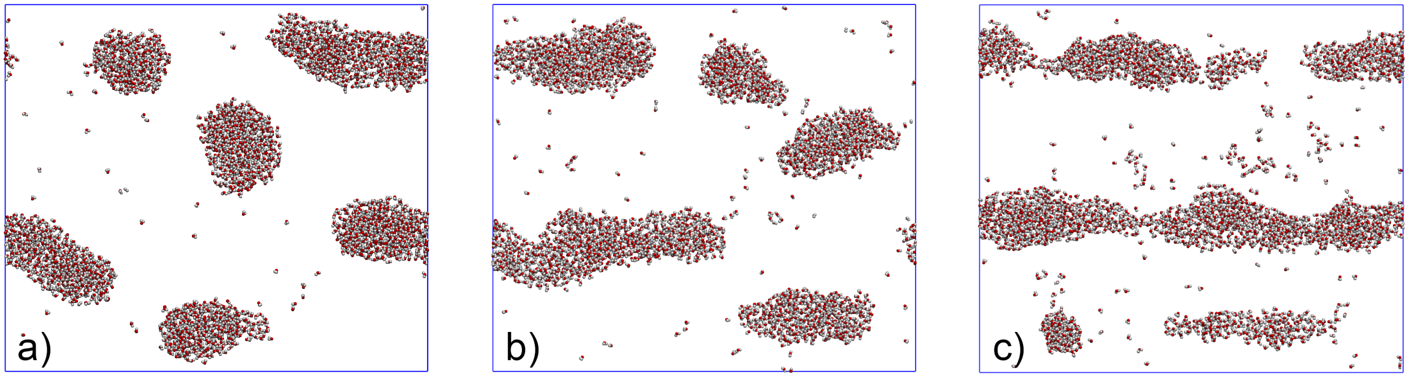


**Figure S7** Top view snapshots of water structures confined between fully-functionalised hair surfaces ($\chi_{\mathrm{N}} = 0$) at sliding speeds of a) $v_s = 0.01$ m s$^{-1}$, b) $v_s = 0.1$ m s$^{-1}$, and c) $v_s = 1$ m s$^{-1}$.

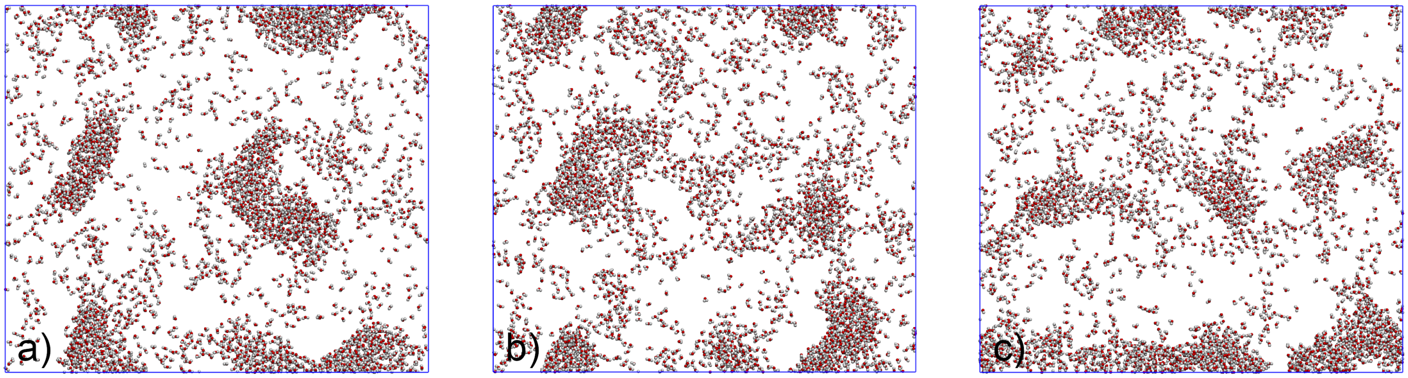


**Figure S8** Top view snapshots of water structures confined between hair surfaces representative of virgin hair ($\chi_{\mathrm{N}} = 0.25$) at sliding speeds of a) $v_s = 0.01$ m s$^{-1}$, b) $v_s = 0.1$ m s$^{-1}$, and c) $v_s = 1$ m s$^{-1}$.

### 7 Wet lipid tilt angles

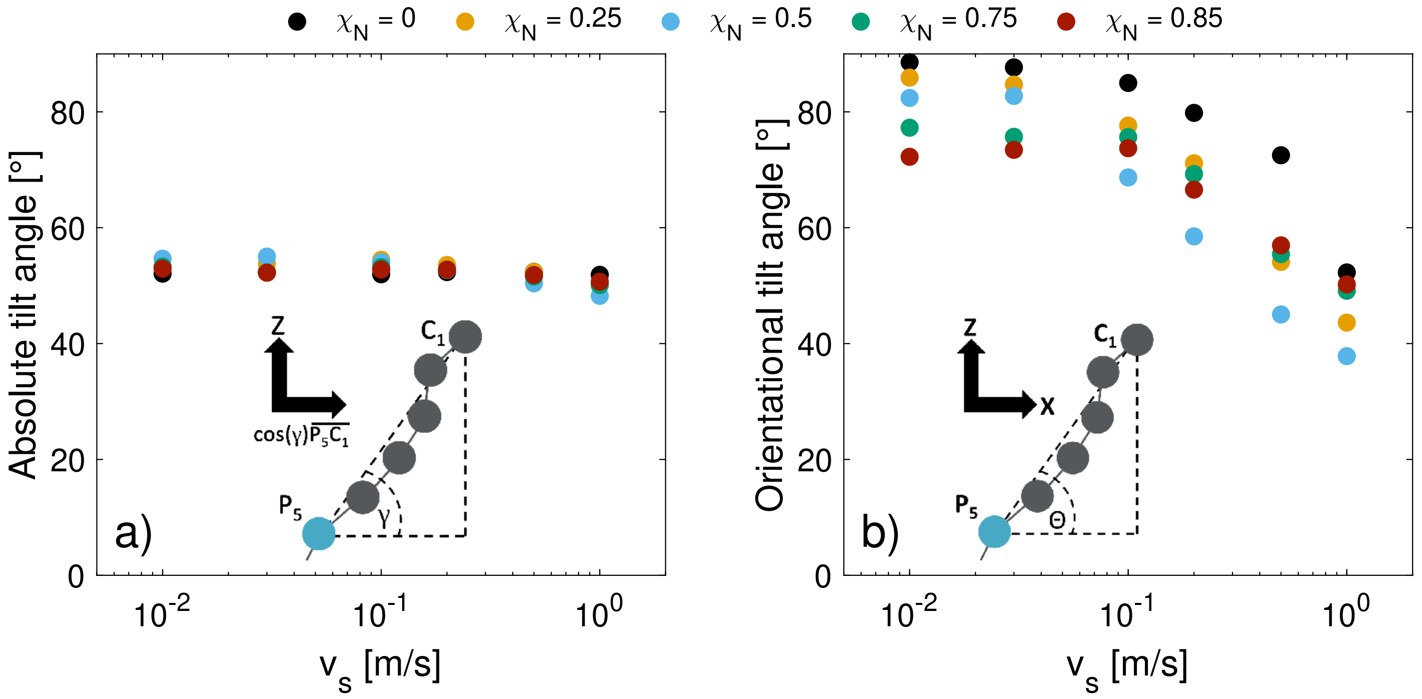


**Figure S9** Absolute lipid tilt angles and orientational tilt angles relative to the direction of sliding for wet contacts as a function of sliding speed and surface damage.

### 8 Wet contact SATA fitting parameters

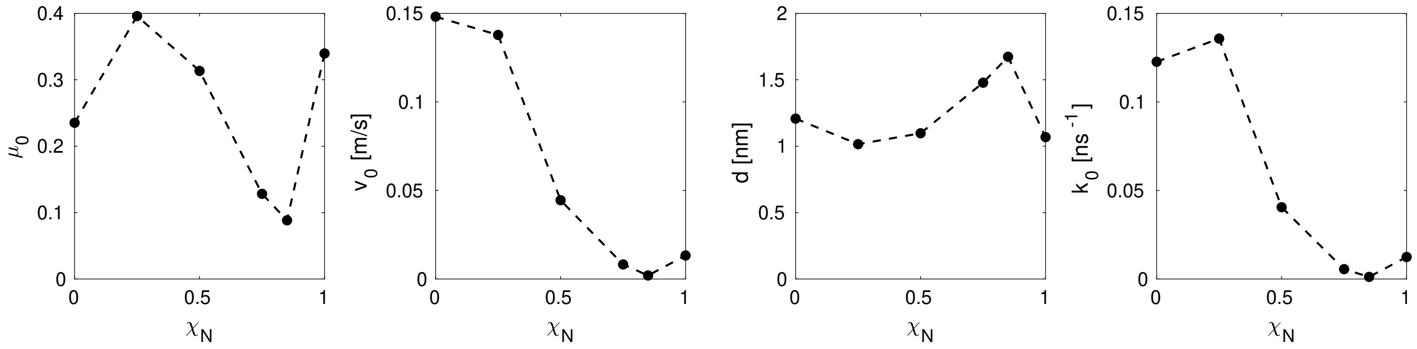


**Figure S10** Wet friction fitting parameters according to Ref.[16] a) Reference CoF $\mu_0$, b) reference velocity $v_0$ and corresponding c) energy barrier distance $d$ and d) hopping rate constant $k_0$.

### 9 Water interfacial area

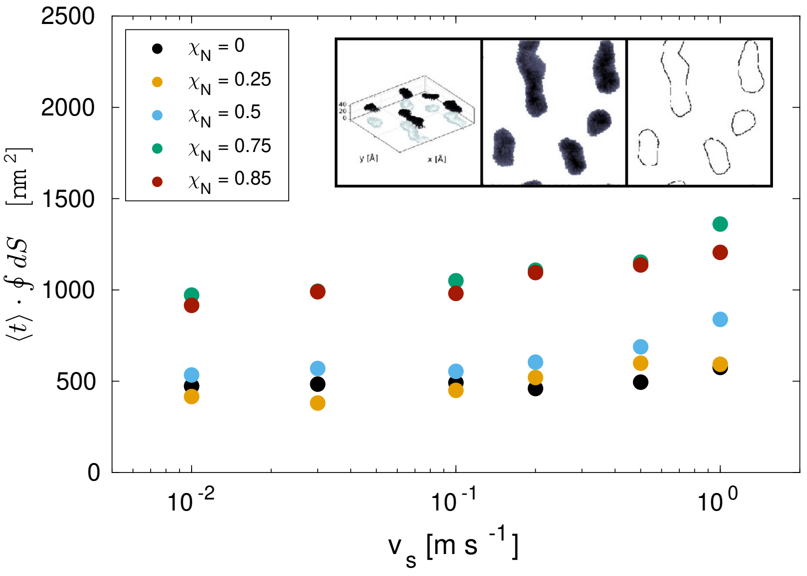


**Figure S11** Water-lipid interfacial area, approximated as the perimeter of each water droplet penetrating the contact down to the thioester bond of the 18-MEA/cysteic acid residues multiplied by the thickness of such structures. The inset shows the extraction of the perimeter by 1) identifying penetrating droplets, 2) obtaining the surface coverage and 3) applying an edge-finding algorithm.

### 10 Counterion structure and diffusion

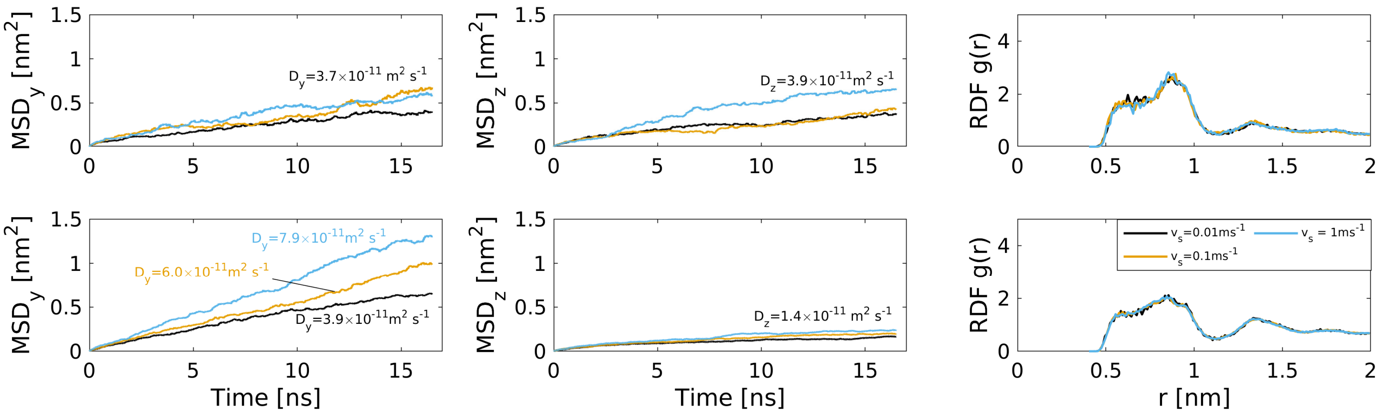


**Figure S12** Counterion mobility by means of MSD$_y$ and MSD$_z$ and RDF as a function of speed. Virgin hair, i.e. $\chi_\mathrm{N} = 0.25$ (top) and medium bleached, i.e. $\chi_\mathrm{N} = 0.85$ (bottom) NEMD model surface configurations are considered.

### 11 Biomimetic friction forces as a function of CCP load

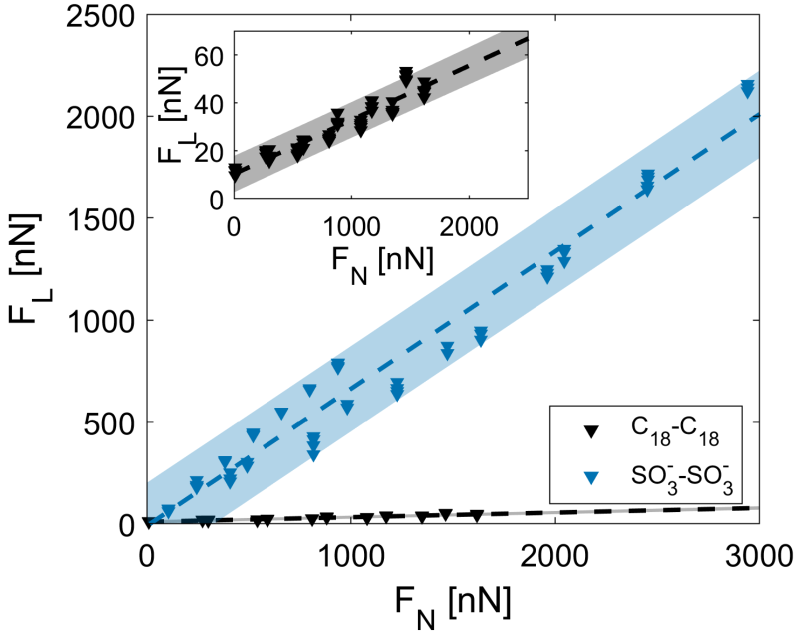


**Figure S13** CCP AFM friction forces as a function of normal force for biomimetic surfaces in water. Linear fits to the raw data are shown as dashed lines. Corresponding prediction intervals on the experimental data with 95% confidence are shown as shaded areas.
